# Inference of multiple mergers while dating a pathogen phylogeny

**DOI:** 10.1101/2023.09.12.557403

**Authors:** David Helekal, Jere Koskela, Xavier Didelot

## Abstract

The vast majority of pathogen phylogenetic studies do not consider the possibility of multiple merger events being present, where a single node of the tree leads to more than two descendent branches. These events are however likely to occur when studying a relatively small population or if there is high variability in the reproductive chances. Here we consider the problem of detecting the presence of multiple mergers in the context of dating a phylogeny, that is determining the date of each of the nodes. We use the Lambda-coalescent theory as a modelling framework and show how Bayesian inference can be efficiently performed using a Billera-Holmes-Vogtmann space embedding and a customised Markov Chain Monte Carlo sampling scheme. We applied this new analysis methodology to a large number of simulated datasets to show that it is possible to infer if and when multiple merger events occurred, and that the phylogenetic dating is improved as a result of taking this information into account. We also analysed real datasets of *Vibrio cholerae* and *Mycobacterium tuberculosis* to demonstrate the relevance of our approach to real pathogen evolutionary epidemiology. We have implemented our new methodology in a R package which is freely available at https://github.com/dhelekal/MMCTime.

## INTRODUCTION

Dated phylogenies have risen to prominence in many research areas of the life sciences, from the study of evolutionary histories of higher organisms, genomic epidemiology of infectious disease, through to understanding diversity of microbial organisms. Most existing approaches to reconstructing and analysing dated phylogenies are restricted to binary trees, where each internal node has exactly two descendent branches. Indeed, Kingman’s coalescent (Kingman 1982) on a continuous real time scale (Drummond et al. 2002) is the most popular framework for modelling dated phylogenies of measurably evolving populations (Drummond et al. 2003; Biek et al. 2015), and in this model only two lineages may merge into the same ancestor at once. However, this model is only applicable if both the sample size and typical family sizes are small in comparison to the effective population size, which can be orders of magnitude smaller than the census population size for example due to heterogeneity of the reproduction success (Charlesworth 2009).

In contrast with the standard Kingman’s coalescent, Lambda-coalescent models, also known as multiple merger coalescents, can be used to describe dated phylogenies where more than two lineages may coalesce into the same ancestor at once (Pitman 1999; Sagitov 1999; Donnelly and Kurtz 1999). Multiple merger events can be the result of various biological phenomena of interest, such as populations undergoing rapid adaptation (Neher and Hallatschek 2013; Desai et al. 2013), superspreading (Hoscheit and Pybus 2019; Lemieux et al. 2020; Gómez-Carballa et al. 2021) or some other form of sweepstakes reproduction (Menardo et al. 2021; Árnason et al. 2023). In particular, the Beta-coalescent (Schweinsberg 2003) is a specific type of Lambda-coalescent that has been used to explain the shallow genealogies observed in cod (Birkner et al. 2013), to study pathogen superspreading (Hoscheit and Pybus 2019) and to characterise *Mycobacterium tuberculosis* outbreak genealogies (Menardo et al. 2021).

Let us take as our starting point an unrooted, undated tree produced by a maximum likelihood tree reconstruction software such as RAxML (Stamatakis 2014), IQ-TREE (Minh et al. 2020) or PhyML (Guindon et al. 2010). Such a tree may contain polytomies, where a node leads to more than two branches. This can be either because of a multiple merger event, or because of at least one branch covering a short interval of time, so that no substitution occurred as expected under any molecular clock model (Bromham and Penny 2003). Multiple heuristic approaches have been developed for either breaking up or collapsing polytomies in undated phylogenies (Kuhn et al. 2011; Lin et al. 2011; Lewis et al. 2005). Here instead we use a Lambda-coalescent framework to infer which polytomies are caused by multiple merger events and which are caused by a lack of phylogenetic signal. We do so in the context of dating the tree, which means to use it as well as the dates of the leaves in order to produce a dated phylogeny (To et al. 2016; Volz and Frost 2017; Didelot et al. 2018). The dated phylogeny can then be used for a broad range of epidemiological investigations (Didelot and Parkhill 2022). To reconstruct this dated phylogeny, we must infer the root position, ancestral node times, as well as parameters associated with the clock and genealogical models. We must distinguish which polytomies are consistent with multiple mergers, and which are better explained by quick binary branching and therefore should be resolved. In the latter case we must also estimate the branching order within the polytomies returned by the maximum likelihood estimation software. This is important as the branching order within the polytomies is random and likely inconsistent with the temporal structure of the tree, as previously noted (Sagulenko et al. 2018).

To achieve these aims several issues must be addressed. We need to choose a set of prior models for the latent genealogies which take into account biological realism and statistical tractability. We focus on the Beta-coalescent (Schweinsberg 2003) and an extension of it described by Eldon and Stephan (2023) in which the Beta-coalescent and Kingman’s coalescent are mixed together, combining low-variance family size reproduction with occasional high-variance sweepstakes. We also need to specify a molecular clock model to establish the relationship between dated and undated phylogenies, and for this we use the Additive Relaxed Clock (ARC) model (Didelot et al. 2021). Next, we need to specify a computational scheme for representing multiple merger trees where a single node may have an arbitrary number of descendants. This representation needs to enable efficient computation of likelihoods and to be statistically efficient. To this end we use the Billera-Holmes-Vogtmann (BHV) space (Billera et al. 2001) augmented with a spike-and-slab construction (George and McCulloch 1993). Finally we design a Markov Chain Monte-Carlo (MCMC) sampling scheme targeting the posterior in order to perform Bayesian inference.

## METHODS

### Lambda-coalescent

Lambda-coalescents are a class of stochastic genealogical processes (Pitman 1999; Sagitov 1999; Donnelly and Kurtz 1999) that generalise the popular Kingman’s coalescent (Kingman 1982) to a setting where more than two lineages may merge into the same parent, i.e. they permit multiple mergers. These processes commonly describe genealogies arising from various forwards in time models in population genetics, typically in scenarios where there is a high variability in the number of surviving offspring, or when selection and recombination are taken into account (Berestycki 2009; Tellier and Lemaire 2014). Examples of such scenarios include heavy-tailed offspring distributions (Schweinsberg 2003; Matuszewski et al. 2018), recurrent selective sweeps in presence of recombination (Durrett and Schweinsberg 2005), rapidly adapting populations (Neher and Hallatschek 2013; Desai et al. 2013), as well as strong purifying selection (Cvijović et al. 2018).

Lambda-coalescents are uniquely specified by a finite measure Λ on [0, 1] that governs the merger sizes of the process. Intuitively, one can think of this relationship as sampling a probability *p* ∼ Λ proportional to the density of Λ, selecting with probability *p* each of the lineages currently in the process, and merging all selected lineages. The instantaneous block merger rates *λ*_*b,k*_, that is the rate at which every *k* ≥ 2 lineages merges into one parent when there are *b* lineages in total, are then given by

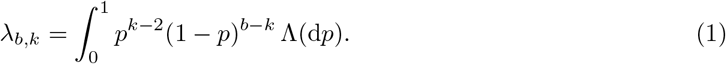

The factor of *p*^−2^ arises from the fact that at least two lineages must participate in order for a merger to happen. When Λ = *δ*_0_, that is a point mass at zero, we can see that

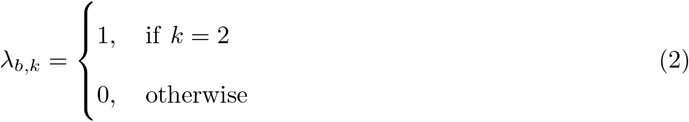

In other words the resulting Lambda-coalescent is Kingman’s coalescent in which only two lineages may merge.

### Beta-coalescent

The approach presented will mostly focus on the Beta-coalescent and an extension of it. The rationale for this is two-fold. First, the Beta-coalescent is relatively well studied, admits the frequently used Kingman’s coalescent as a special case, and the instantaneous block merger rates are available in closed form. Second, it arises from models in which the variance of the offspring distribution can be very high, and has been considered in the context of pathogen populations before, see for example (Hoscheit and Pybus 2019). As the name suggests, in the case of the Beta-coalescent the measure Λ is simply the Beta distribution. Usually in the context of Lambda-coalescents, the Beta distribution is parameterised as (Schweinsberg 2003)

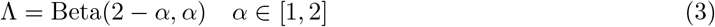

The reason for this is a connection to a model of populations with skewed offspring distributions in which *α* governs the tail behaviour of the offspring distribution. The forwards-in-time model in the derivation of (Schweinsberg 2003) follows dynamics of a supercritical Galton–Watson process where in successive non-overlapping generations each of the *N* individuals produces *v*_*i*_ offspring i.i.d. according to a distribution with the tail index *k*^− *α*^. This implies that the offspring distribution has infinite variance if 1 *< α <* 2 and infinite mean if *α* = 1. Offspring are then randomly killed in order to keep the population size constant and equal to *N* . Schweinsberg (2003) showed that the genealogies arising for this process converge to Kingman’s coalescent for values of *α* ≥ 2 and to the Beta-coalescent parameterised as in Equation 3 for values of *α* ∈ (2, 1], under suitable time-rescalings and as *N* → ∞. In plain words if the distribution of the number of offspring produced by any of the individuals is sufficiently skewed, and this situation arises frequently enough, every once a while an individual may get lucky and produce enough offspring to replace a non-negligible fraction of the population. Therefore the probability that multiple individuals find the same ancestor at once does not vanish in the large population size limit. The parameter *α* relates to how skewed the offspring distribution is. The limiting case of *α* = 1 corresponds to the Bolthausen-Sznitman coalescent (Bertoin and Le Gall 2000), which has been shown to arise in scenarios corresponding to rapid adaptation and clonal interference (Desai et al. 2013; Neher and Hallatschek 2013; Schweinsberg 2017). On the other hand, in the limiting case of *α* = 2, the Beta distribution collapses into an atom at 0 and thus the resulting coalescent is the Kingman’s coalescent (Schweinsberg 2003).

There are several limitations of the Beta-coalescent. First amongst these is the assumption that every individual may produce a large number of offspring in every generation. Often it is easier to imagine that such large reproductive events may only occur if correct circumstances are met. Related to this is the second limitation. The Beta-coalescent has a time scale of *N* ^1− *α*^ if *α* ∈ (1, 2] and log (*N*)^−1^ if *α* = 1. For many populations, this implies mutation rates or population sizes orders of magnitude higher than what would be biologically realistic. This is especially the case if *α* is close to 1 as previously noted (Eldon and Stephan 2023). This may however not be an issue for moderately large values of *α* in the case of within-host evolution where the population sizes are going to be very large.

### Extended Beta-coalescent

We now introduce an extension of the Beta-coalescent which we will refer to as the extended Beta-coalescent. This modification is a mixture of the Beta-coalescent and Kingman’s coalescent. This is achieved by defining the measure Λ characterising this process as

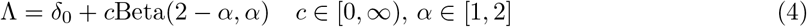

The reasons for introducing this are two-fold. For one it will provide us with a convenient example of a Lambda-coalescent of a form shared by for example the Durrett–Schweinsberg coalescent (Durrett and Schweinsberg 2005) arising from selective sweeps. The second reason is based on the modifications to random sweep-stakes reproduction presented in (Eldon and Stephan 2023). They considered a modification to the construction in (Schweinsberg 2003) to address the problematic assumption of very frequent large family sizes. In this modification, in each generation a coin with probability *E* is flipped. On success with probability *E* each individual produces offspring according to an offspring distribution with a “small” *α* ∈ [1, 2) and with probability 1 − *E* according to an offspring distribution *α* ≥ 2. For a suitable choice of *ϵ* = *ϵ* _*N*_ → 0 as *N* → ∞, the resulting coalescent process is exactly the Lambda-coalescent specified by Equation 4.

### Selection of priors

For the Beta-coalescent we parametrise the measure characterising the genealogical prior as:

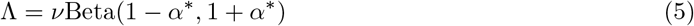

*v* and *α* ^∗^ are unknown parameters we wish to infer. *v* ∈ ℝ_+_ is the time scale of the process and *α* ^∗^ ∈ [0, 1] controls the Beta distribution governing the merger sizes. *α* ^∗^ relates to the original parameter *α* from Equation 3 (Schweinsberg 2003) as *α* = *α* ^∗^ + 1. We equip these parameters with the following prior distributions:

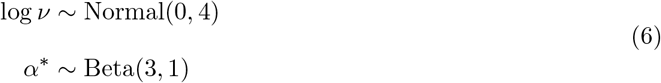

We caution that it is not straightforward to interpret *v* as the timescale of the Beta-coalescent is 1*/N* ^1− *α*^ and therefore *v* only corresponds to the usual effective population size if *α* = 2.

For the extended Beta-coalescent we parameterise the genealogical prior as:

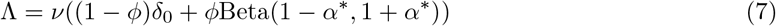

*v* ∈ ℝ_+_ corresponds to the process rate, *ϕ* ∈ [0, 1] is the mixing proportion between the Kingman component and the Beta component, and *α* ^∗^ ∈ [0, 1] once again controls the Beta distribution and therefore the size of multiple mergers. We equip these parameters with the following prior distribution:

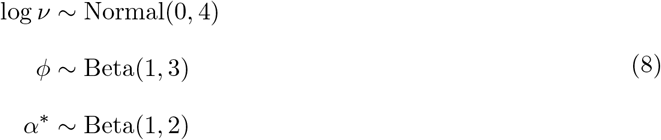

Finally, for the molecular clock model we use the Additive Relaxed Clock (ARC) (Didelot et al. 2021) in which a branch of length *l* carries a number of substitutions *x* distributed as:

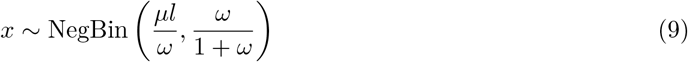

where *μ* represents the mean clock rate and *ω* controls the amount of relaxation relative to a strict clock model. We use the following prior distribution:

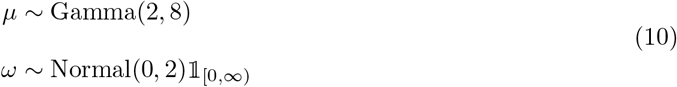

### Billera-Holmes-Vogtmann space embedding

In order to efficiently sample genealogies that admit multiple mergers we leverage two concepts. First we embed binary trees in a space of phylogenetic trees where coordinates correspond to branch lengths, specifically the Billera-Holmes-Vogtmann (BHV) space. Second we use a spike-and-slab construction to put positive mass on the set of trees where at least one branch length is shrunk to exactly zero and identify these trees with multiple merger trees obtained by collapsing all branches with lengths equal to exactly zero.

BHV space is a metric space introduced in the study of phylogenetic tree geometry (Billera et al. 2001). For a fixed number of tips *n* the BHV space is constructed from a set of (2*n* − 3)!! orthants in (0, ∞)^*n*−2^ each corresponding to a particular topology of a rooted binary tip labeled *n*-tree. Each coordinate within the *n* − 2 dimensional orthant corresponds to the length of an interior branch in the given tree topology. At any of the zero boundaries, the binary topology degenerates into a tree that has branching greater than two. The orthants are “glued” together at boundaries corresponding to the same topology. For example if only one coordinate approaches zero there is a junction of three different *n* − 2 dimensional orthants corresponding to an *n* − 3 dimensional orthant face. For existence of centroids, proof of constant negative curvature and properties of geodesics refer to (Billera et al. 2001). We will base our sampling scheme construction around the BHV space. The BHV space has been used previously in phylogenetic reconstruction for example to construct an embedding amenable to sampling binary trees using discrete Hamiltonian Monte Carlo (Dinh et al. 2017). Typically such approaches simply select the base measure to be Lebesgue within each of the *n*−2 dimensional orthants corresponding to binary topologies, and each *n* − 2 dimensional orthant is assigned equal probability. The lower dimensional orthants are then null sets within this measure space. However such an approach is not appropriate for inference with multiple merger trees as any set consisting purely of these would be given a zero probability. One option would be to put mass on lower dimensional orthants, and set up a trans-dimensional sampling scheme using reversible jump MCMC (Green 1995). However this would be unlikely to work efficiently as designing moves that remove or add more than one branch at a time and don’t suffer from a rapidly diminishing acceptance ratio would be challenging, and would not take advantage of the natural geometry of the space.

We will therefore augment the BHV space using a spike-and-slab construction (George and McCulloch 1993). Denote by 𝒯 the set of all rooted, labeled, metric *n*-trees. An *n*-tree is said to be metric if all of its branch lengths are strictly greater than 0. Let 𝕋 denote the set of all labeled, rooted, binary *n*-tree topologies. Denote the closed *n* − 2 dimensional orthant of the BHV space corresponding to a particular binary *n*-tree topology as *V* ^*τ*^ for *τ* ∈ 𝕋. We identify points in the closed *n* − 2 dimensional orthants of BHV space by the tuple (*X, Q, τ*) where *X* ∈ [0, ∞)^*n*−2^ denotes the location within an orthant, *Q* ∈ {0, 1}^*n*−2^ is a vector of indicators where *q*_*i*_ = 1 if and only if *x*_*i*_ = 0 and *τ* ∈ 𝕋 denotes the orthant index, i.e. the corresponding binary topology. We next identify all points on the boundary *∂V* ^*τ*^, that is all points with at least one coordinate equal to zero with the corresponding *k*-ary metric *n*-tree by collapsing each and every branch for which *q*_*i*_ = 1, i.e. that is of length exactly zero. We now define the base measure to be

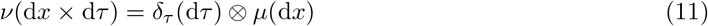

That is we assign uniform mass to each of the *n* − 2 dimensional orthants and within each orthant we have

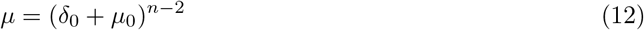

Where *δ*_0_ is an atom at zero and *μ*_0_ is the Lebesgue measure on [0, ∞). This construction assigns positive probability to sets of binary trees with one or more branches with length exactly zero. By identifying such trees with a (metric) multiple merger tree we can see that it therefore puts a positive probability on trees with multiple mergers.

### Parametrisation of the genealogy

Having outlined the construction above we now give an explicit parametrisation of the genealogy that will allow us to construct an MCMC scheme as well as enable the computation of necessary quantities. As an input we assume we are given a rooted binary phylogeny with *n* tips, with branch lengths corresponding to the estimated number of substitutions along a branch. This phylogeny is assumed to be a point estimate obtained by ML phylogenetic software. The root position may be assumed to be known a-priori, or to be estimated, in which case the initial rooting is assumed to be chosen arbitrarily. In practice the estimated number of substitutions along a branch may not be an integer even though it is likely to be very close to one when the number of substitutions per site is low. The clock models used require an integer number of mutations. Therefore all branch lengths are coerced to integer values by rounding. Based on the undated input phylogeny we can construct a rooted binary genealogy denoted as ***τ*** as follows. We define the following notations:

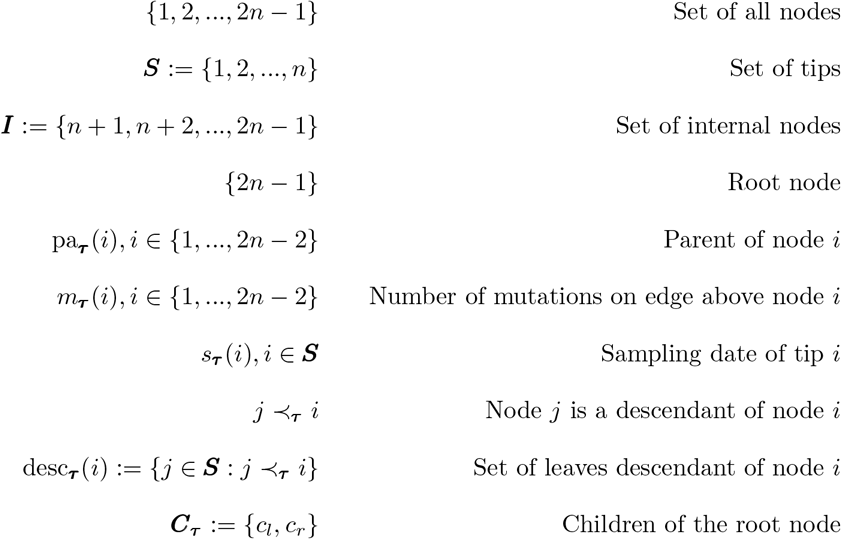

Having defined necessary notation for characterising the rooted tree topology we aim to define the parametrise the internal node heights in the rooted genealogy. This corresponds to the within orthant parametrisation. We now proceed to define the collection of node times ***t*** := {*t*_*i*_}_1≤*i*≤2*n*−1_. A subset of these variables corresponds to the internal node times, that is the free parameters of the model. We denote these as ***t***^***I***^ := {*t*_*i*_ ∈ ***t*** : *i* ∈ ***I***}. The remaining first *n* variables correspond to sampling times and these are considered as an input. Denote these set of sampling times by ***t***^***S***^ := {*t*_*i*_ ∈ ***t*** : *i* ∈ ***S***}. In practice, the vector ***t*** needs to satisfy a complex set of constraints, that is

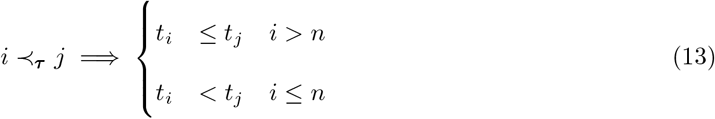

Where *i* ≺_***τ***_ *j* denotes the partial order induced by the tree topology, that is *i* ≺_***τ***_ *j* iff *i* is a descendant of *j*.

The complex set of constraints that the node times have to satisfy would make defining a Metropolis-Hastings move capable of targeting the boundaries complicated. Hence we define the vector of positive height variables 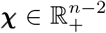 as well as the tree height variable *H* ∈ ℝ. Once again for convenience we will abuse notation and denote by χ (*i*) the component of **χ** associated with node *i*. This is in contrast to *χ*_*i*_ which denotes the *i*-th component of the vector **χ**. To define the transformation for **χ** to ***t*** we first need to introduce the set of lower bounds *b*_***τ***_

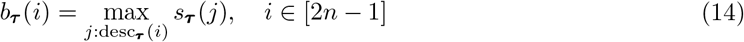

We now map χ, *H* to ***t*** via the mapping *g* : (*H*, ***χ***) ⟼ ***t***. The first part of the mapping parametrises the height of the root node by transforming *H*

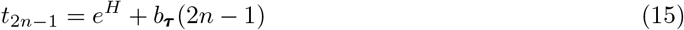

The second part of mapping parametrises the height of internal non-root nodes by transforming ***χ***

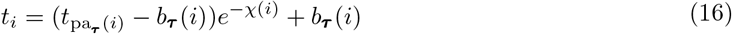

Note that *χ* (*i*) = 0 implies that the length of the branch above node *i* scaled in time units is also 0. An illustration of how such parametrisation looks in practice can be found in Figure 1.

**Figure 1:**
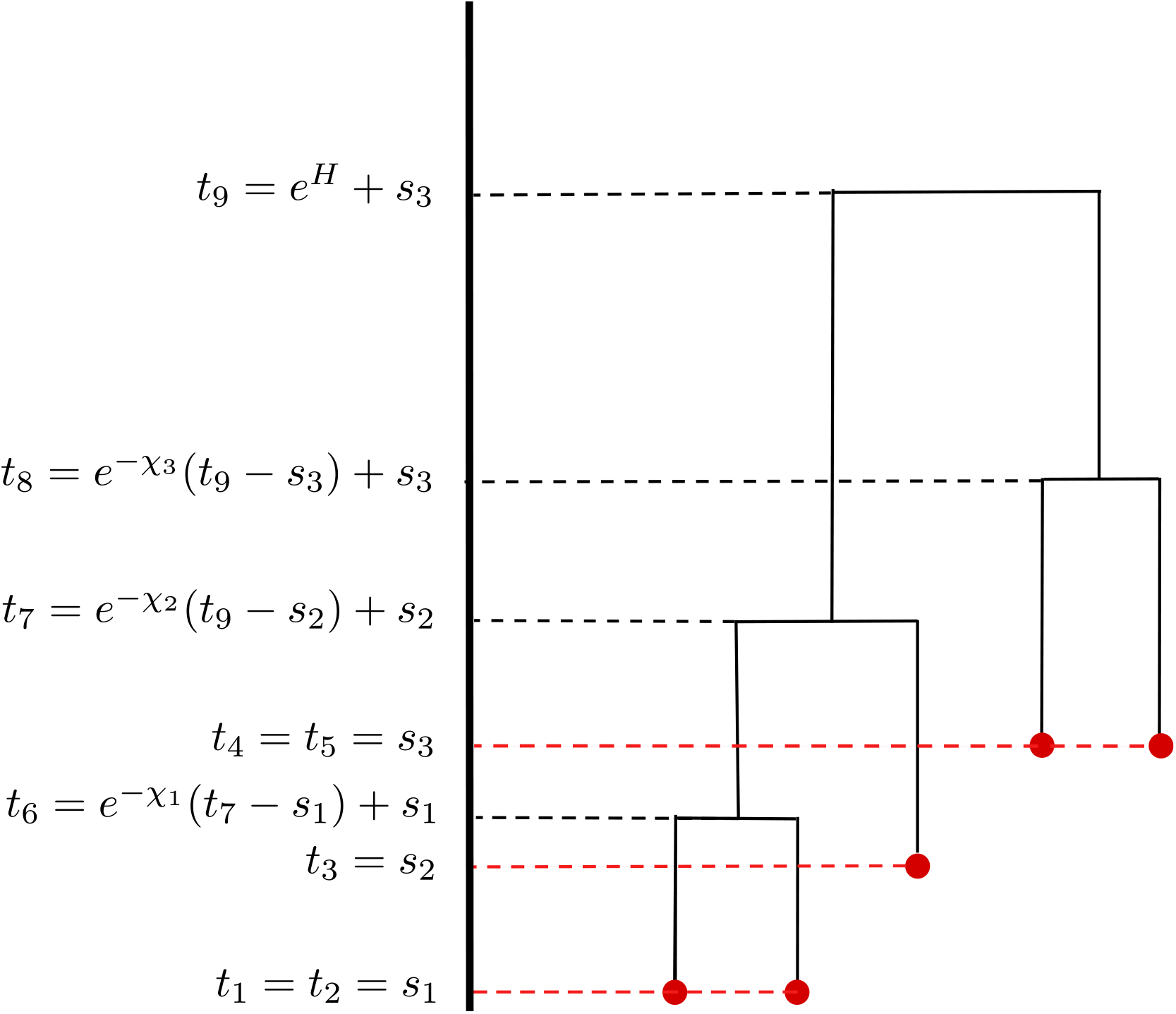
Example of tree parametrisation. In this case the bounds are *b*(6) = *s*_1_, *b*(7) = *s*_2_, *b*(8) = *s*_3_, *b*(9) = *s*_3_. The components ***χ*** are mapped as *χ* (6) = *χ* _1_, *χ* (7) = *χ* _2_, *χ* (8) = *χ* _3_.

This parametrisation is reminiscent of the ratio transform of (Ji et al. 2021), with the key difference that we express non-root internal node heights in terms of a transform of the distance from the parent constrained so that the age of the child is strictly less than the oldest descendant tip as opposed to a transform of the ratio of the remaining height. This is because it is necessary in the construction presented here for the boundary corresponding to 0 branch length to be accessible. Finally we introduce the vector of indicators that determine whether a coordinate is allocated to the boundary:

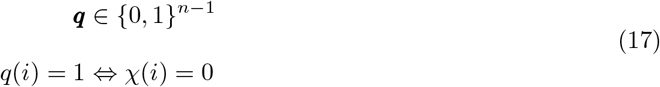

We can now parametrise a genealogy by the tuple ***Y*** := (*H, χ*, ***q***, *τ* ; ***s***) where *τ* specifies the binary topology and hence a full dimensional orthant of the BHV space, and *H, χ*, ***q*** characterise the position within the orthant conditional on the tip date constraints ***s***.

### Lambda-coalescent likelihood

In order to compute the coalescent likelihood of a genealogy, we first need to convert the (possibly non-metric) binary tree embedding to the respective metric multiple merger tree. To achieve this first denote the set of all internal nodes descending from node *i* as

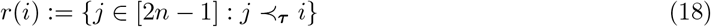

Next defining the set of internal descendants of *i* coincident with *i* as

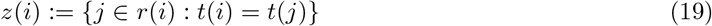

We can see that for any internal node *i* such that *q*(*i*) = 0, |*z*(*i*)| corresponds to the merger size minus 2. Using these to definitions we can compute the likelihood of the binary tree embedding ***Y*** := (*H, χ*, ***q***, *τ* ; ***s***) under the Lambda-coalescent Λ^*θ*^ with block merger rates 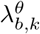 as:

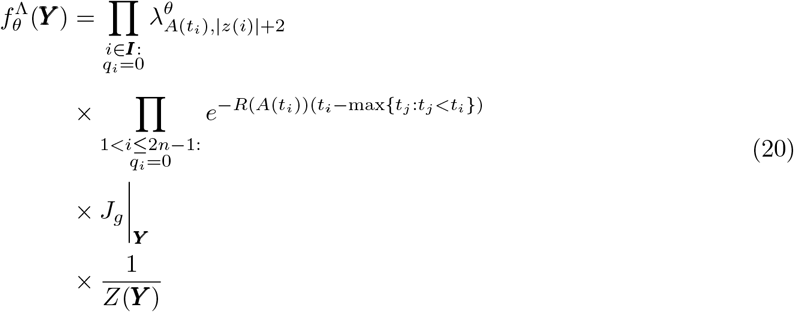

Where *R* : ℕ ⟼ ℝ_+_ denotes the total coalescent rate of the process

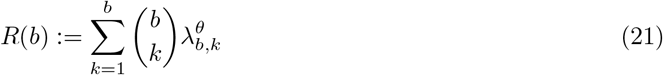

and *A* : ℝ_+_ ℕ _+_ denotes the lineages through time function at time *t*, defined as

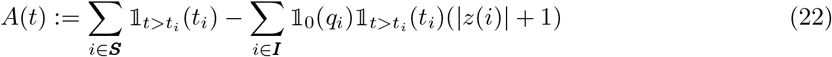

This is also known as the block counting process (Kukla and Möhle 2018). As the genealogical prior is expressed in terms of node heights ***t***, we must account for the transformation from ***χ*** to ***t*** in the density. To do so we require the Jacobian for the mapping *g* : (*H*, ***χ***) *1*→ ***t***. This is straightforward to compute as the matrix of first order partials has a diagonal structure and hence is equal to

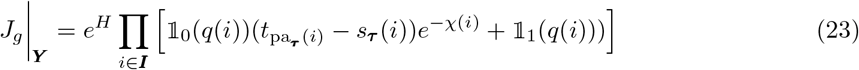

Note that as the density of all *χ* _*i*_ = 0 is with respect to an atomic measure these do not play a role within the relevant Jacobian adjustment. Finally we note that the embedding of a given multimerger tree as a binary tree is not unique. Therefore, in order for the density of a given tree to be proportional to the density given by the Lambda-coalescent density, we need to re-weight the density of the embedding to account for the overcounting. In general given a multiple merger of size *m* there are (2*m* − 3)!! ways to resolve this as a sequence of binary mergers. This is the number of rooted labeled binary trees with *m* tips (Billera et al. 2001). Therefore the adjustment for the embedding *Y* is:

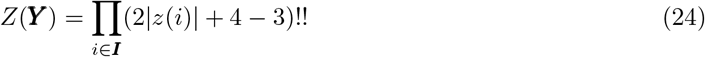

### Branch likelihood

The likelihood is based on the ARC model (Didelot et al. 2021). Each branch in the input ML phylogeny is scaled in estimated number of substitutions along that branch. This number of substitutions is distributed according to Equation 9. The number of substitutions returned by ML software will in general not be an integer, although in many cases will be close to an integer value. As the ARC model requires integer valued number of substitutions all values are rounded to the nearest integer. Define the length of a branch above node *i* as 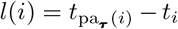. Given the current genealogy state parameterised by ***Y***, and the clock parameters *μ* and *ω*, the likelihood for all branches not incident to the root can be computed as:

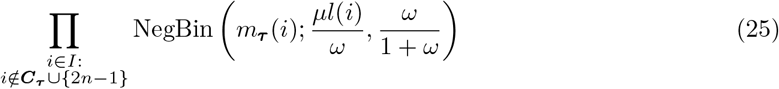

This has to be multiplied by the likelihood of the branches incident to the root. This depends on whether the root is considered fixed, or if it is unknown. If the root is fixed, the branches incident to root are treated as any other branch and their contribution is

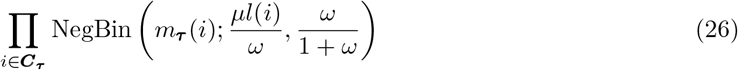

If the root position is considered to be unknown then the position of the root on the branch is marginalised out and the contribution becomes

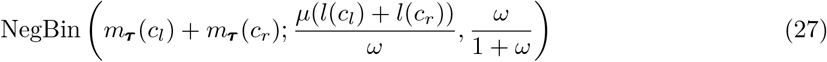

This follows from the additive nature of the ARC clock model (Didelot et al. 2021).

### MCMC scheme

We use four types of moves to sample parameters characterising *Y* . These moves can be categorised into three families. The first family consists of moves covering transitions that update the position within a full dimensional orthant including the boundary. The moves within the first family update branch lengths, and therefore merger sizes and internal node heights. The second family consists of moves for proposing transitions between full dimensional orthants. The moves within the second family update the root position and branching order within polytomies. The third family consists of a single move that updates the parameters of the observation model and genealogy model. The scheme consists of a single sweep through all three families, selecting one move from each family uniformly at random.

### Orthant interior move

The orthant interior move is a random walk Metropolis (RWM) (Metropolis et al. 1953) move restricted to updating a subset of the vector ***χ*** corresponding to those coordinates which are currently not restricted to the boundary, and thus a part of a multiple merger

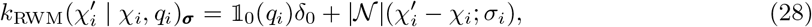

where | 𝒩 |(*·*; *σ*) denotes the density of the modulus of a normal random variable with variance *σ*^2^. As expected for RWM the proposal ratio equals to one.

### Orthant boundary move

The orthant boundary move is responsible for transitioning between binary topologies and multiple mergers. To do so the move needs to propose transitions between the boundary and orthant interior. The main challenge with designing this move is that due to the structure of the likelihood of the extended Beta-coalescent there is a relatively sharp ridge between a 2-merger and a 3-merger when the number of active lineages is large, i.e. *b* » 3. Therefore the move must be able to propose transitions that move several coordinates of *χ* to 0 or back at once. The first step consists of sampling an internal node index *i* ∈ ***I*** | {2*n* − 1} at random with probability

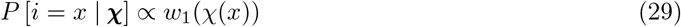

If the coordinate is allocated to the interior (therefore to the slab component of the base measure), i.e. if *q*(*i*) = 0, *χ*(*i*) *>* 0 then the move begins by proposing *χ*^*t*^(*i*) = 0. Denoting the parent node above *j* = pa(*i*) the moves continues upwards and proposing to shrink *χ*(*j*) to 0 with probability *w*_2_(*χ*(*x*)) as long as it is accessible to this move, i.e. *q*(pa(*j*)) = 0 and *j* ≠ 2*n* − 1. This process repeats until a coordinate fails to shrink either due to it not being accessible or due to the coin flip failing.

If the coordinate initially selected is allocated to the boundary then there are two options. The move either expands that coordinate, proposing *χ*^*t*^(*i*) *>* 0. Alternatively the move expands that coordinate and then attempts to expand the coordinate above it if it is allocated to the boundary. If the coordinate above it is allocated to the boundary it is expanded and this procedure repeats, terminating when either the root is reached or a non-zero coordinate is reached. The expanded coordinates are sampled from a proposal distribution *d*, i.e. *χ*^*t*^ ∼ *d*. Whether the move expands coordinates recursively or not is decided uniformly at random. Denote the sequence of nodes that have had their corresponding coordinates modified by *v*_1_, *v*_2_, …, *v*_*m*_, where *v*_1_ is the first node modified by the move. If the initial coordinate chosen was allocated to the interior the proposal ratio is equal to

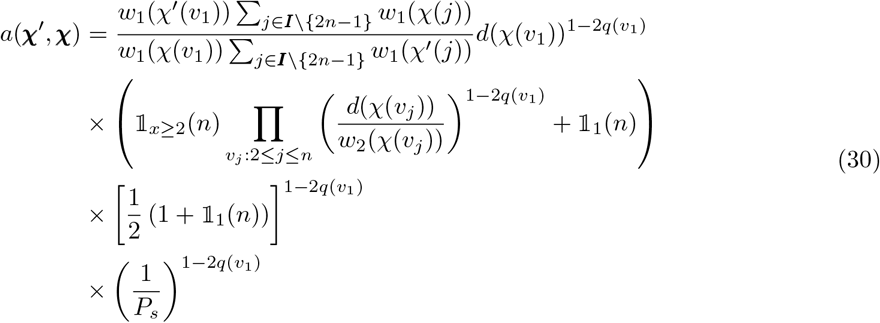

Note that 1 − 2*q*(*v*_1_) = 1 if the move is shrinking coordinates towards the boundary and −1 if it is expanding coordinates. Therefore it determines the direction of the move. The first term corresponds to the probability of selecting the same starting node, the second term corresponds to the likelihood of the coordinate transformation for subsequent nodes, the third accounts for the possibility of the reverse move being chosen, and the fourth term corresponds accounts for the stopping probability of the recursion, *P*_*s*_ which is equal to

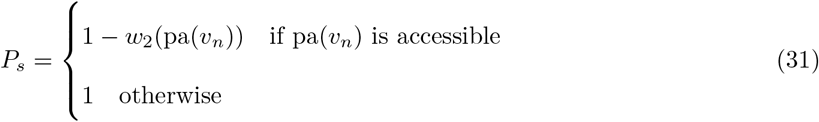

Crucially setting *w*_2_(χ) to be equal to the density of *d*(χ) leads to the second term cancelling to one. In practice we use

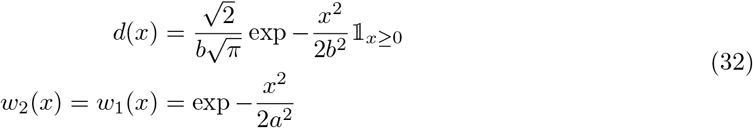

Setting 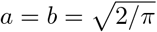 leads to *d*(*x*) = *w*_1_ (*x*) = *w*_2_(*x*).

### Root NNI move

The root NNI move updates the root position, and is used if the root position is considered unknown. It is a version of the nearest neighbour interchange (NNI) commonly used in phylogenetic inference (Yang and Rannala 1997). This moves first proceeds in selecting one of the child nodes of the root as a pivot. The move then moves the root to one of the descendant edges of the pivot. The pivot is chosen with equal probability from both root descendant nodes at random with the exception of two special cases. If one of the descendant nodes is a tip node the other descendant of the root is selected as a pivot. This is because NNI move is undefined for a tip node selected as a pivot. The other special case is if one of the root descendants has the branch above it collapsed as a part of a multiple merger but the other does not. In this case the descendant with the edge collapsed is chosen as the pivot. This is to prevent the move from wasting computational effort as this way the node with edge collapsed stays adjacent to the root. Otherwise it may become adjacent to an edge that cannot support a multiple merger leading to 0 likelihood. With the pivot selected the branch to move the root to is sampled with equal probability from two edges that descend from the pivot. We denote the pivot by *p* ∈ ***C***_***τ***_ and its probability mass function under the topology ***τ***

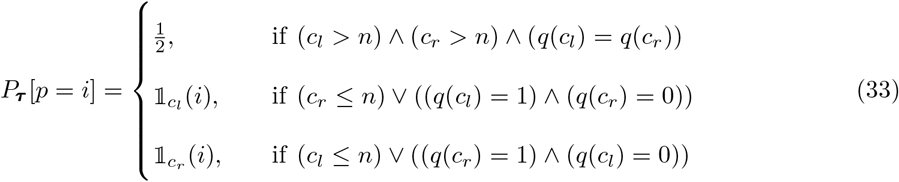

With *p* fixed, sample *j* from the two descendants of *p* uniformly at random. Based on this generate a new rooted tree topology ***τ*** ^*t*^, which is the same as ***τ*** except where:

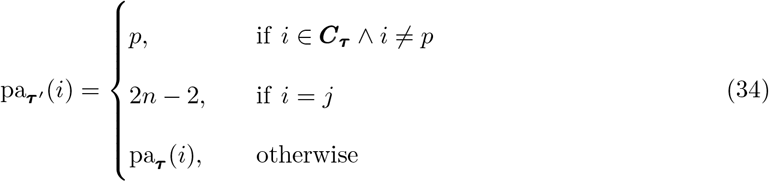

The mutations above the affected nodes are then adjusted accordingly:

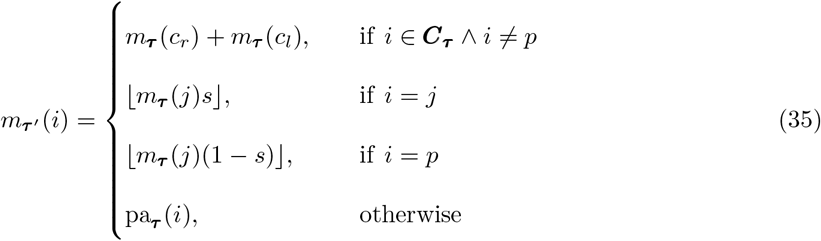

For some arbitrarily chosen *s* ∈ [0, 1].

The proposal ratio is

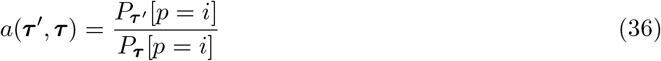

### Polytomy neighbour interchange move

The polytomy neighbour interchange move changes the binary topology within polytomies as this topology is randomly resolved (if at all) by the ML estimation program. It does so by first sampling a pivot node *p* from all nodes for which the edge above contains 0 mutations. With the pivot selected a node *j* is chosen uniformly at random from all the descendants of *p* that are adjacent to the polytomy. That is from all descendant nodes such that the parent of that node is 0 mutations away from *p*. Denote this set of nodes *A*_***τ***_ . Next a node *k* is selected from all nodes in the outgroup relative to *p* such that the distance of the parent of those nodes to the parent of *p* is 0 mutations. Denote this set of nodes *B*_***τ***_ . Based on this generate a new rooted tree topology ***τ*** ^*t*^, which is the same as ***τ*** except where:

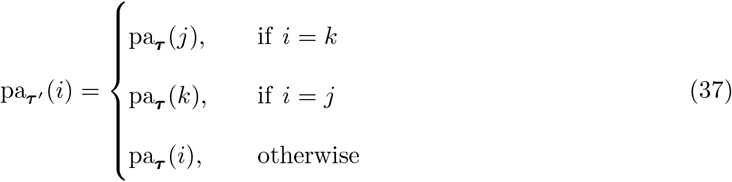

The reverse move consists of selecting the same pivot and then the corresponding descendants. Denote the sibling of the pivot *p* with *s*_*p*_. The proposal ratio is

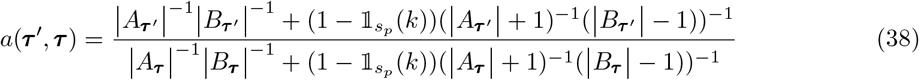

### Tuning the sampler

In order to tune the sampler a preconditioning matrix of individual parameter variances is estimated, along with a scale factor multiplying this matrix. The RWM moves updating the parameters and node height variables are then scaled accordingly. This is done in a sequence of six steps. Let *T* denote the thinning factor. The first two steps consists of running the sampler with orthant boundary moves disabled. The first step consists of burn-in for 100 *× T* iterations. The second step then estimates the *χ* preconditioning matrix for 100 *× T* iterations. For the following steps all moves are enabled. The third step consists of burn-in for 100 *× T* steps. The fourth step estimates the preconditioning matrix for the parameter moves for 100 *× T* steps. The fifth and sixth steps estimate the step scaling for 50 *× T* steps. All previous iterations are then discarded.

## RESULTS

### Implementation

We implemented the approach as described in an new R package titled *MMCTime*, which is available at https://github.com/dhelekal/MMCTime. The package uses *ape* (Paradis and Schliep 2019) as a backend for handling phylogenies. *bayesplot* is used for handling MCMC diagnostic visualisations, (Gabry et al. 2019) and *ggtree* (Yu et al. 2017) is used for visualising phylogenies. The package *posterior* (Vehtari et al. 2021) is used for computing MCMC diagnostics.

### Illustration on simulated phylogenies

All simulations follow the same protocol. First a dated phylogeny is simulated from a genealogical model, conditional on appropriate parameter values and tip sample times. Then the expected number of substitutions for each branch is sampled using the ARC model (Didelot et al. 2021). Then Seq-Gen (Rambaut and Grass 1997) is used to generate sequences of a given length under the HKY model (Hasegawa et al. 1985). For all simulation experiments the length was set to 10000bp, except if otherwise mentioned. Finally, a ML phylogenetic tree is reconstructed using IQ-TREE (Minh et al. 2020). This, along with the tip dates, serves as a starting point for the analysis. For all simulation benchmarks, the root position was assumed to be unknown and to be inferred.

A first dated phylogeny was simulated under the Beta-coalescent (Equation 5) with parameters {*v* = 1*/*12, *α*^∗^ = 1*/*2} as shown in Figure 2A. The ARC clock model with parameters {*μ* = 1, *ω* = 1} was applied, and the input ML phylogeny is shown in Figure S1. Inference was performed under the Beta-coalescent model using four chains, sampling every 2000 iterations for a total of 1000 samples retained per chain. Assessing mixing and qualities of estimates is challenging in this setting as the topology changes due to the uncertain branching on polytomies. Furthermore standard metrics like the Robinson-Foulds distance (Robinson and Foulds 1981) are inappropriate for dated trees and not applicable to multiple merger trees. To circumvent this we compute effective sample sizes (ESS) and 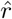 estimates for the genealogical parameters as well as the clock parameters along with the tree height and the two following summaries: the number of multiple mergers and the maximum merger size. The 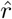 and ESS estimates are computed using technique in (Vehtari et al. 2021) as implemented in the R package `posterior`. This confirmed that the MCMC had converged and mixed as expected. The MCMC traces are shown in Figure S2 and the inferred parameters in Figure S3, with all inferred ranges covering the correct values. Nine posterior samples of the dated phylogeny are shown in Figure S4. To summarise the full posterior sample of dated phylogenies, we use a modified version of the DensiTree representation (Bouckaert 2010) as shown in Figure 2B. Comparison of the simulated (Figure 2A) and inferred (Figure 2B) phylogenies demonstrate the accuracy of the inference, including the identification of which nodes are likely to be multiple merger events.

**Figure 2:**
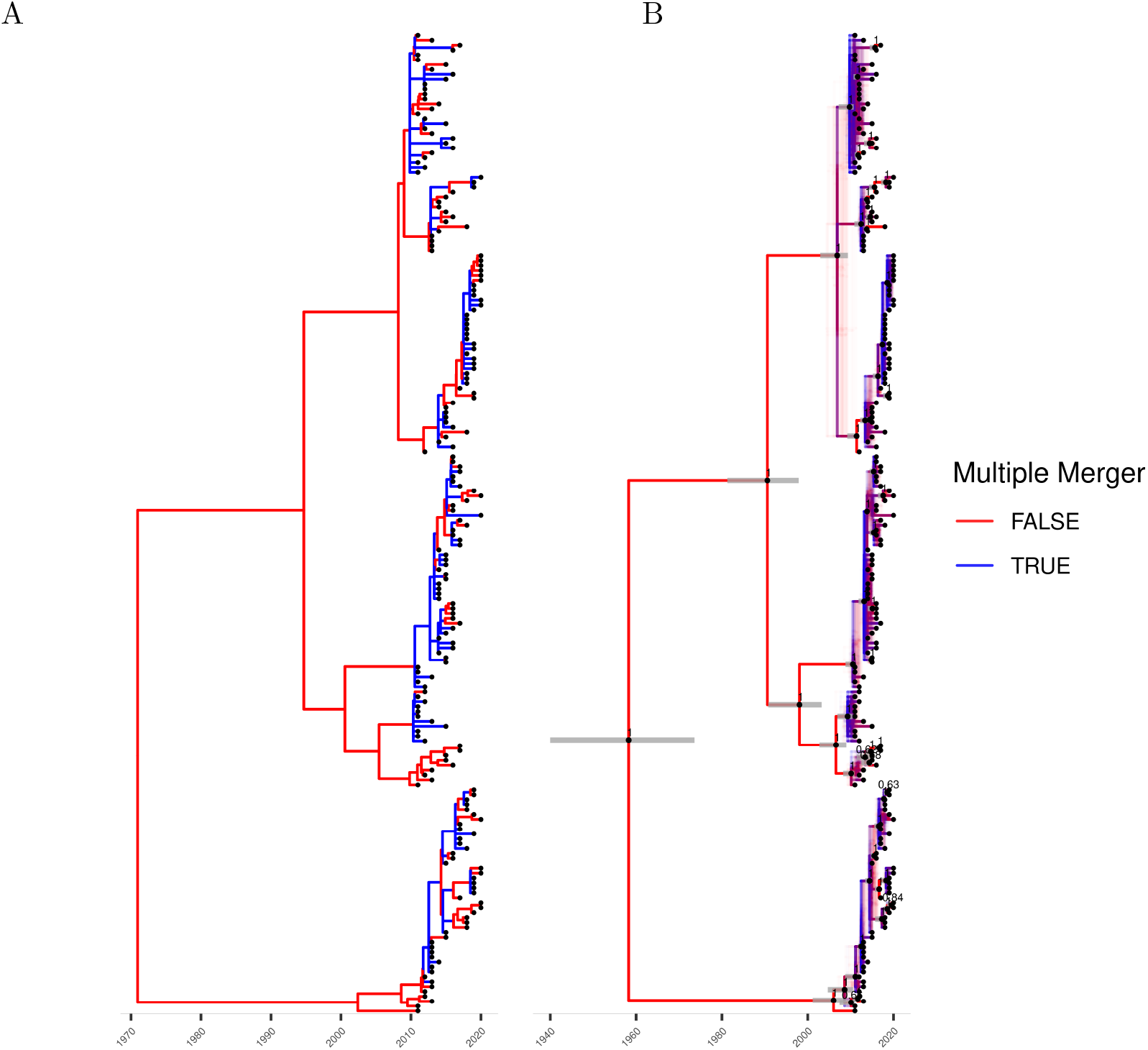
Simulation and inference under the Beta-coalescent model. (A) The simulated phylogeny. (B) A qualitative summary of the posterior, showing locations possible multiple mergers and uncertainty in polytomy topology. Clades appearing in over 50% of posterior samples are indicated with black dots fixed at median height, and grey bars overlayed indicated the 95% posterior credible interval for the height of these nodes.

Next a dated phylogeny was simulated under the extended Beta-coalescent (Equation 7) with parameters {*v* = 1*/*12, *α*^∗^ = 1*/*5, *ϕ* = 2*/*5} (Figure 3A). The same analysis as above was performed, except that the extended Beta-coalescent model was used for inference. The ML tree is shown in Figure S5, the MCMC traces in Figure S6, the parameters in Figure S7, nine posterior sampled phylogenies in Figure S8 and the posterior phylogeny summary in Figure 3B. Once again we find that the parameters and phylogeny are inferred satisfactorily. There were only a few multiple merger events in the simulated tree, most of which behaved as a Kingman’s coalescent tree. This represents a good illustration of what can be achieved with the extended Beta-coalescent, and would be very unlikely to happen under the Beta-coalescent model.

**Figure 3:**
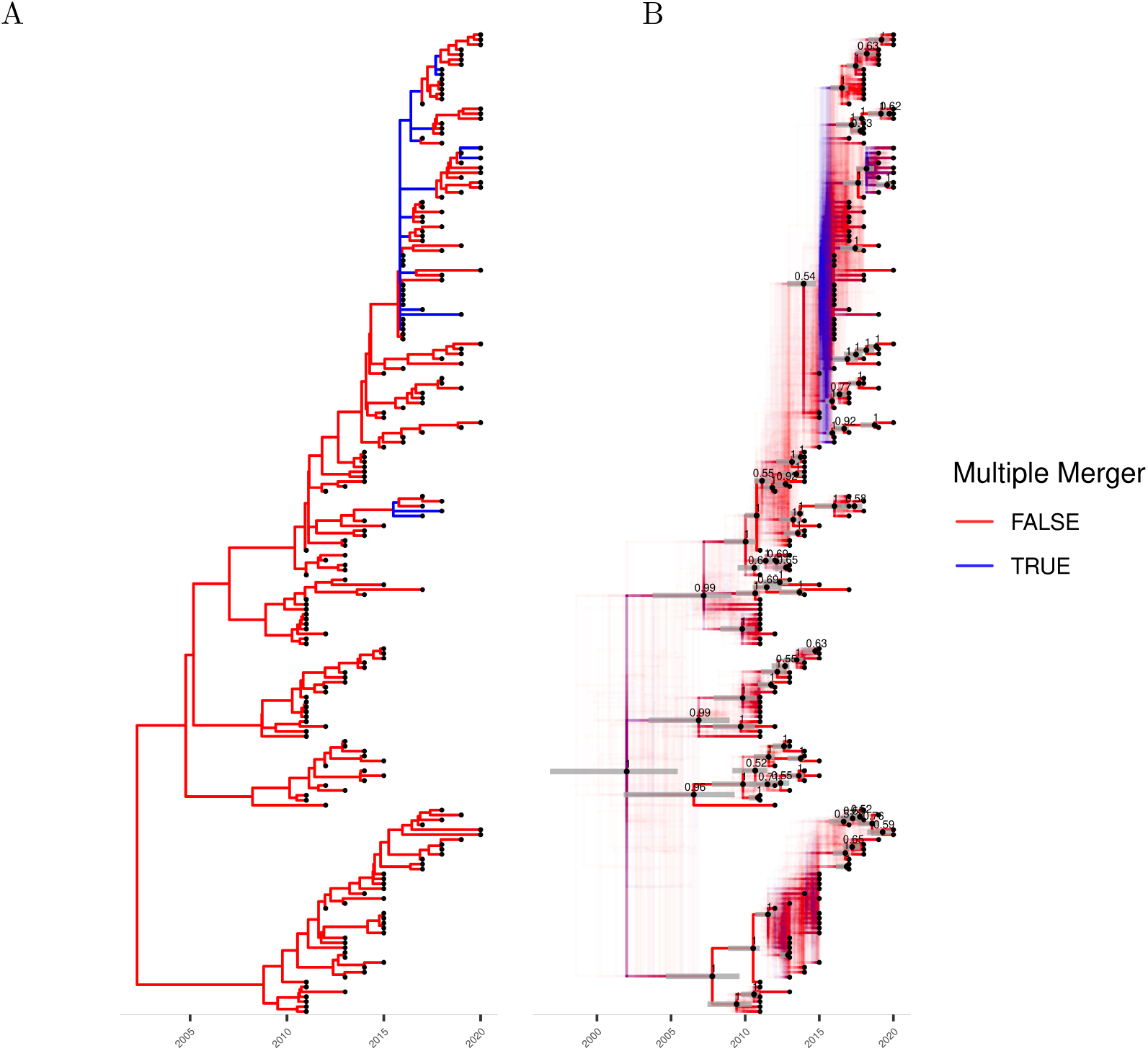
Simulation and inference under the extended Beta-coalescent. (A) The simulated phylogeny. (B) A qualitative summary of the posterior, showing locations possible multiple mergers and uncertainty in polytomy topology. Clades appearing in over 50% of posterior samples are indicated with black dots fixed at median height, and grey bars overlayed indicated the 95% posterior credible interval for the height of these nodes.

### Benchmark under the Beta-coalescent

To benchmark the performance of inference under the Beta-coalescent, we considered three ARC clocks with the following parameters: Clock 1 has *μ* = 1.5 and *ω* = 0.5, Clock 2 has *μ* = 3 and *ω* = 1.0, Clock 3 has *μ* = 6 and *ω* = 2. Figure S9 shows the distributions of the number of substitutions per site under each of these clock models. For each clock model, Figure 4 shows the results of parameter inference under the Beta-coalescent for 150 datasets generated with *v* = 0.1 and values of *α*^∗^ increasing between 0 and 1. In every case the parameters are correctly inferred, except for *α*^∗^ which is slightly overestimated when the correct value was lower than 0.1. Low values of *α*^∗^ lead to trees with a high probability of large multiple merger events. To avoid this biologically implausible scenario we used a Beta(3,1) prior for inference, which only has cumulative probability 0.001 for *α*^∗^ ∈ [0, 0.1]. The use of this prior explains the slight overestimation of *α*^∗^ when the correct value was low.

**Figure 4:**
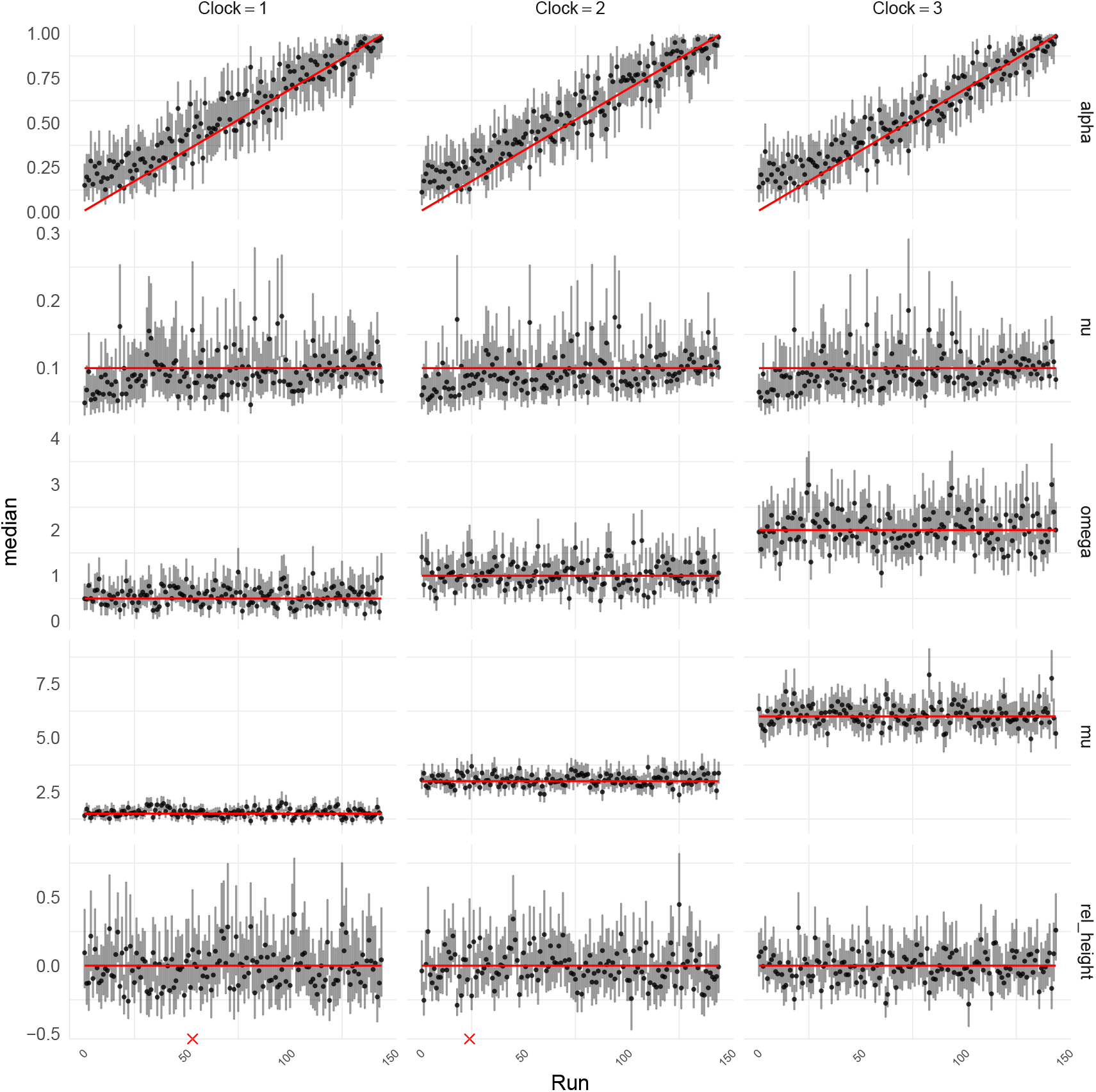
Simulation benchmark under the Beta-coalescent. Posterior summaries for analysis of limitations of the likelihood approximation. Red lines indicate ground truth. Vertical bars represent 95% posterior credible intervals, and points represent the median. Red crosses indicate insufficient mixing in the corresponding.

Figure S10 shows the number of nodes in the inferred phylogenies minus the true number of nodes in the simulated phylogenies. Positive values indicate an excess of nodes and therefore an underestimation of the number of multiple merger events. Negative values indicate a lack of nodes and therefore an underestimation of the number of multiple merger events. Most intervals cover the correct value of zero. There is a slight tendency to overestimate the number of nodes overall, which is again driven by our use of a conservative prior for *α*^∗^.

### Benchmark under the extended Beta-coalescent

We performed a similar benchmark for the inference under the extended Beta-coalescent. Four scenarios were considered with *φ ∈ {*0, 0.5, 0.75, 1*}*. Each scenario consists of 48 genealogies. Within a scenario the values of *α*^*\**^ were linearly varied from *α*^*\**^ = 0.01 to *α*^*\**^ = 0.75. The upper limit was chosen to 0.75 since as *α*^*\**^ *→* 1 the process behaves like Kingman’s coalescent irrespective of the value of *φ*, rendering the scenario meaningless. As for the previous benchmark the genealogies were then used to generate three datasets each with different clock parameters. Figure S11 shows the distributions of the number of substitutions per site under each of the three clock models.

The inferred values of the parameters *φ* and *α*^*\**^ are shown in Figures S12 and S13, respectively. The scenario *φ* = 0 corresponds to the Kingman’s coalescent. In this scenario the parameter *φ* was consistently estimated to be low as expected, and *α*^*\**^ could not be estimated (i.e. the posterior was approximately equal to the prior) since in this scenario this parameter does not play a role. In the scenarios where the Kingman’s coalescent and Beta-coalescent were mixed with *φ* = 0.5 and *φ* = 0.75 the parameter *φ* was usually underestimated to the point that the inferred values of *α*^*\**^ did not follow the correct values. However, in the scenario *φ* = 1, which corresponds to the pure Beta-coalescent, it was possible to infer the values of *φ* and *α*^*\**^ as long as *α*^*\**^ was not too high. When *α*^*\**^ is high the Beta-coalescent component of the extended Beta-coalescent prior behaves like the Kingman’s coalescent component, so that the mixing proportion *φ* does not have much effect on the data.

Thus the extended Beta-coalescent suffers from identifiability issues on the parameters *φ* and *α*^*\**^ in the part of the parameter space where the model reduces to the Kingman’s coalescent, namely when *φ* is low and/or *α*^*\**^ is high. This does not affect the estimates of the remaining parameters though. The parameters *ν, μ* and *ω* are shown in Figures S14, S15 and S16, respectively, and are all well estimated. Figures S17 and S18 show that the time to the most recent common ancestor and number of nodes in the tree are also estimated around their correct values. Note that in the scenario with *φ* = 0 the correct tree is completely binary and so the number of nodes can only be underestimated. Finally, Figure S19 shows the estimated probabilities that a tree sampled from the posterior contains a multiple merger, which increases as expected as *φ* increases.

### High mutation rate limitation

A difficulty arises when the mutation rate per site is too high. In this case the probability of reversal or homoplasious mutation increases, such that the maximum likelihood estimated branch lengths of the input phylogeny become unlikely to be exactly zero even when a multiple merger event occurred. This is related to the branch saturation observed in dating methods that use a maximum likelihood tree as input, such as LSD (To et al. 2016) or BactDating (Didelot et al. 2018). In order to gain an understanding of when this phenomenon becomes problematic, we repeated the simulation benchmark under the Beta-coalescent but with a genome length of 1000bp (i.e. 10 times less than previously) and mutation rates doubled for each clock model, i.e. *μ ∈ {*3, 6, 12*}*. Figure 5 shows the results for this analysis. There is a clear and consistent overestimation of *α*^*\**^ when the correct value of this parameter was low. This corresponds to bias against configurations of the Beta-coalescent that produce larger multiple mergers. This was accompanied by underestimation of the process timescale parameter *ν*. This bias worsens as the mutation rate increases and thus the expected number of substitutions per site increases. This result shows that the approach presented here is not appropriate when the number of substitutions per site is too high, specifically in the order of *μ ≈* 10 per year. Since the simulated trees had sums of branch lengths in excess of 100 years (eg Figure 4A), and the number of sites simulated was 1000bp, this corresponds to an expected number of substitutions greater than one for each site, which would not be an issue in practice for the applications envisaged here.

**Figure 5:**
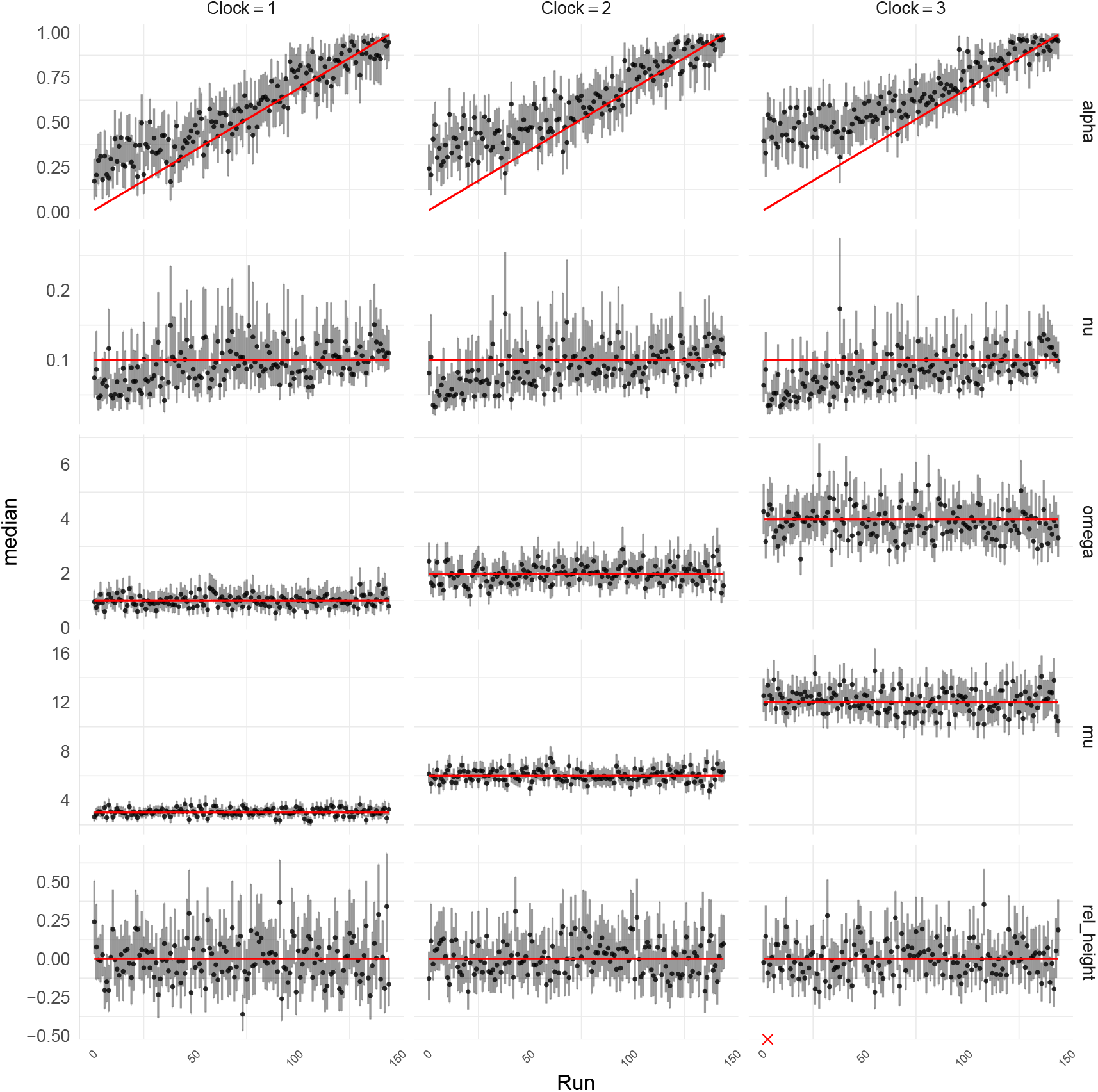
Posterior summaries for analysis of limitations of the likelihood approximation. Red lines indicate ground truth. Vertical bars represent 95% posterior credible intervals, and points represent the median. Red crosses indicate insufficient mixing in the corresponding run.

### Case Study: Spread of *Vibrio cholerae* in Argentina

A recent study compared genome sequences of *Vibrio cholerae*, the causative agent of cholera, sampled from Argentina and neighboring countries between 1992 and 2000 in order to characterise its population structure (Dorman et al. 2020). We selected from the previously published phylogeny the genomes that had been isolated in Argentina and for which the isolation date was known, resulting in a phylogeny containing 411 leaves as shown in Figure S20. We applied inference under the extended Beta-coalescent model, which produced the traces shown in Figure S21 and the parameter estimates shown in Figure S22. The rooting of the tree was fixed using an outgroup. Nine samples from the posterior phylogeny are shown in Figure S23. A phylogenetic posterior sample is summarised as a DensiTree in Figure 6A. This contains several large well supported multiple merger events, consistent with the high estimate of *φ* and low estimate of *α*^*\**^.

**Figure 6:**
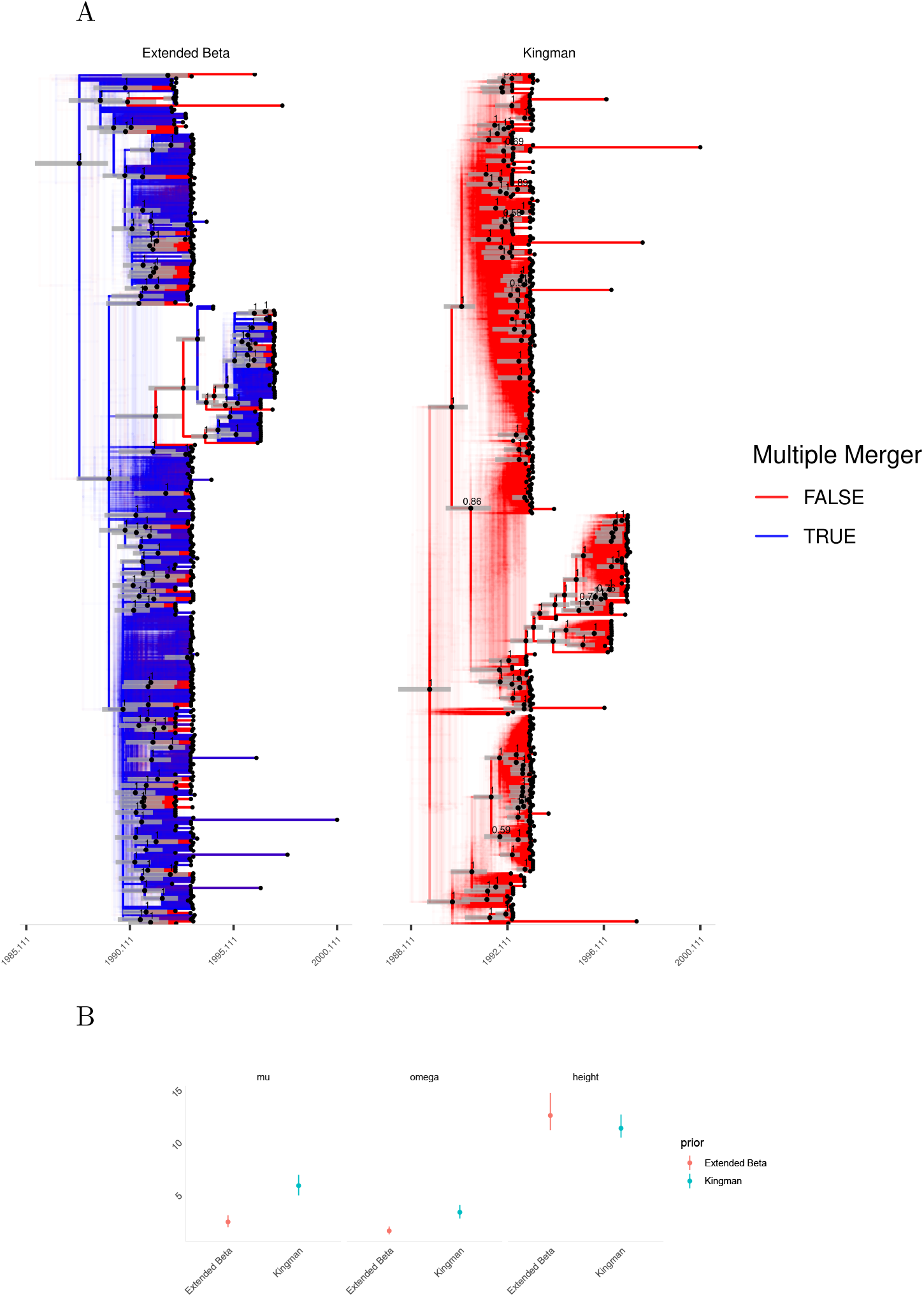
Analysis of the *Vibrio cholerae* dataset. (A) Qualitative summaries of the posterior inferred under the extended Beta-coalescent and Kingman’s coalescent, showing locations possible multiple mergers and uncertainty in polytomy topology. Note that the tip ordering is not identical between the two summaries. Clades appearing in over 50% of posterior samples are indicated with black dots fixed at median height, and grey bars overlayed indicated the 95% posterior credible interval for the height of these nodes. (B) A comparison of estimated parameters under the extended Beta-coalescent and Kingman’s coalescent models.

For comparison purposes, we also performed inference under the pure Kingman’s coalescent model, and nine samples from the posterior phylogeny are shown in Figure S24. As we can see in Figure 6B the clock rate estimated for the *Vibrio cholerae* genealogy under Kingman’s coalescent is much higher than the one under the extended Beta-coalescent. Furthermore the relaxation parameter *ω* is higher when using the Kingman’s coalescent, indicating that evolution is less clock-like. The estimated mutation rate under the extended Beta-coalescent falls into a posterior 95% credible interval of [1.90 - 2.84] mutations per genome per year. This is in good agreement with previous estimates of the *V. cholerae* clock rate based on sparsely sampled worldwide collections of genomes (Mutreja et al. 2011; Didelot et al. 2015). In contrast the substitution rate estimated under Kingman’s coalescent is higher with credible interval [4.98 - 6.64] mutations per genome per year, which is inconsistent with previous estimates. Consequently, the time to the most recent common ancestor for the whole Argentinian dataset is underestimated when using Kingman’s coalescent as opposed to the extended Beta-coalescent (Figure 6B).

### Case Study: *Mycobacterium tuberculosis* outbreak phylogenies

The importance of multiple merger genealogies to study tuberculosis outbreaks has been recently demonstrated (Menardo et al. 2021) using data from eleven previously published outbreaks. We selected three of these for reanalysis, labelled *Bainomugisa2018* (Bainomugisa et al. 2018), *Eldholm2015* (Eldholm et al. 2015) *and Lee2015* (Lee et al. 2015). *These three datasets were selected because they had more than 90% probability of the model selected being a Beta-coalescent in the previous analysis (Menardo et al. 2021) and their phylogenies were not being excessively large. Analysis was performed for each of the three phylogenies under three models: the extended Beta-coalescent, the Beta-coalescent and the Kingman’s coalescent. The three input trees are shown in Figure S25. The rooting of the trees was fixed using outgroup rooting*.

*As can be seen in Figure 7, analysis under Kingman’s coalescent leads to a higher clock relaxation parameter ω* value, suggesting that this model is less appropriate. This is also shown by the fact that *α*^*\**^ was always inferred much smaller than one. The effect is most pronounced for the *Eldhom2015* dataset. In the extended Beta-coalescent the parameter *φ* was estimated to be very close to one for this dataset, in which case it becomes approximately equivalent to the Beta-coalescent. A qualitative summary of the genealogies inferred for this dataset can be seen in Figure 8. Nine realisations of the dated genealogies are shown for the Beta-coalescent, extended Beta-coalescent and Kingman’s coalescent in Figures S26, S27 and S28, respectively. As expected, the results under the Beta-coalescent and extended Beta-coalescent are very similar, including evidence for several large multiple merger events. These results are in good agreement with the previous analysis of these three datasets (Menardo et al. 2021), and highlight the importance of considering multiple mergers when analysing phylogenetic data from tuberculosis outbreaks.

**Figure 7:**
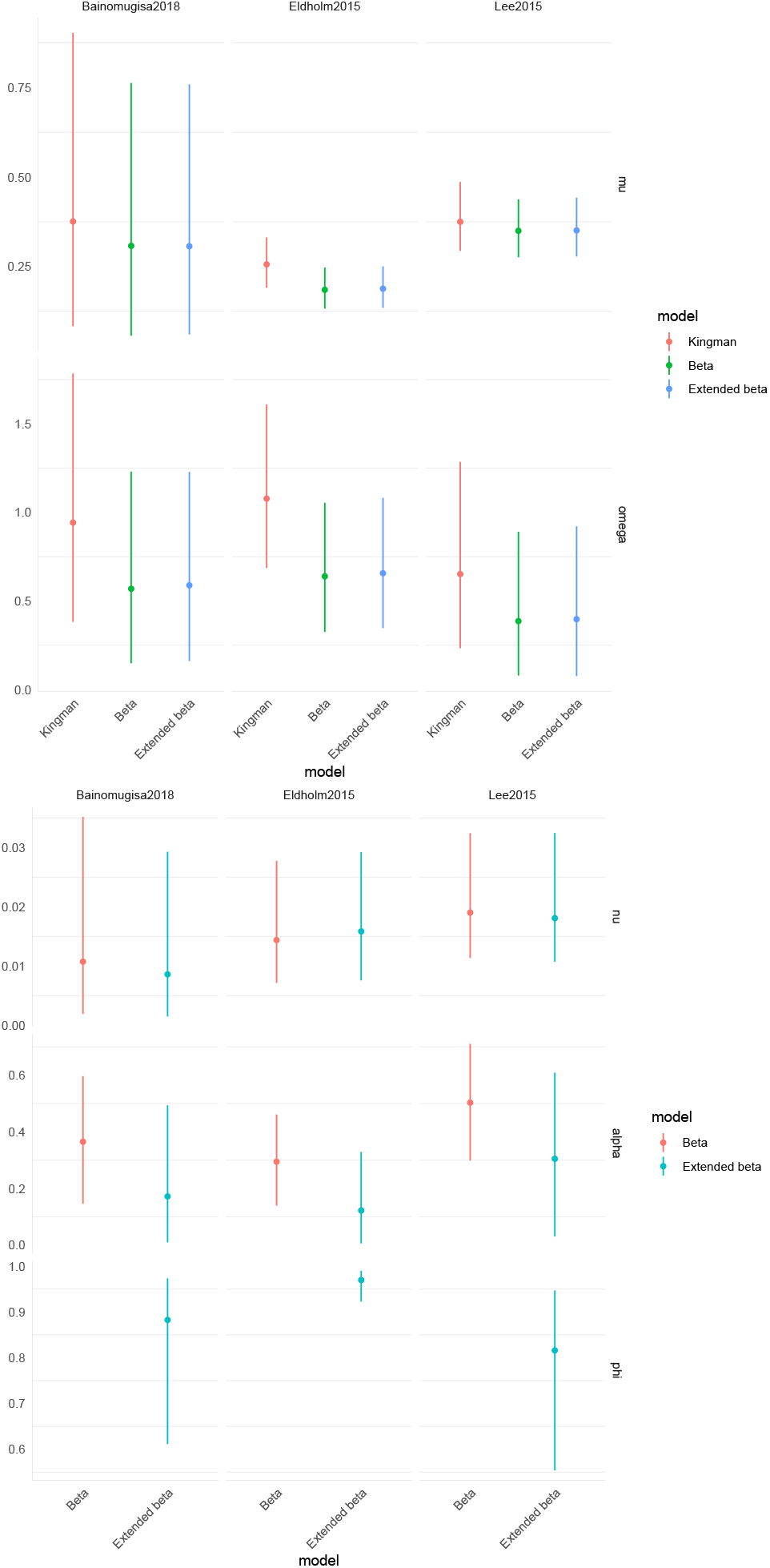
A comparison of parameter marginals estimated under different models for each dataset.

**Figure 8:**
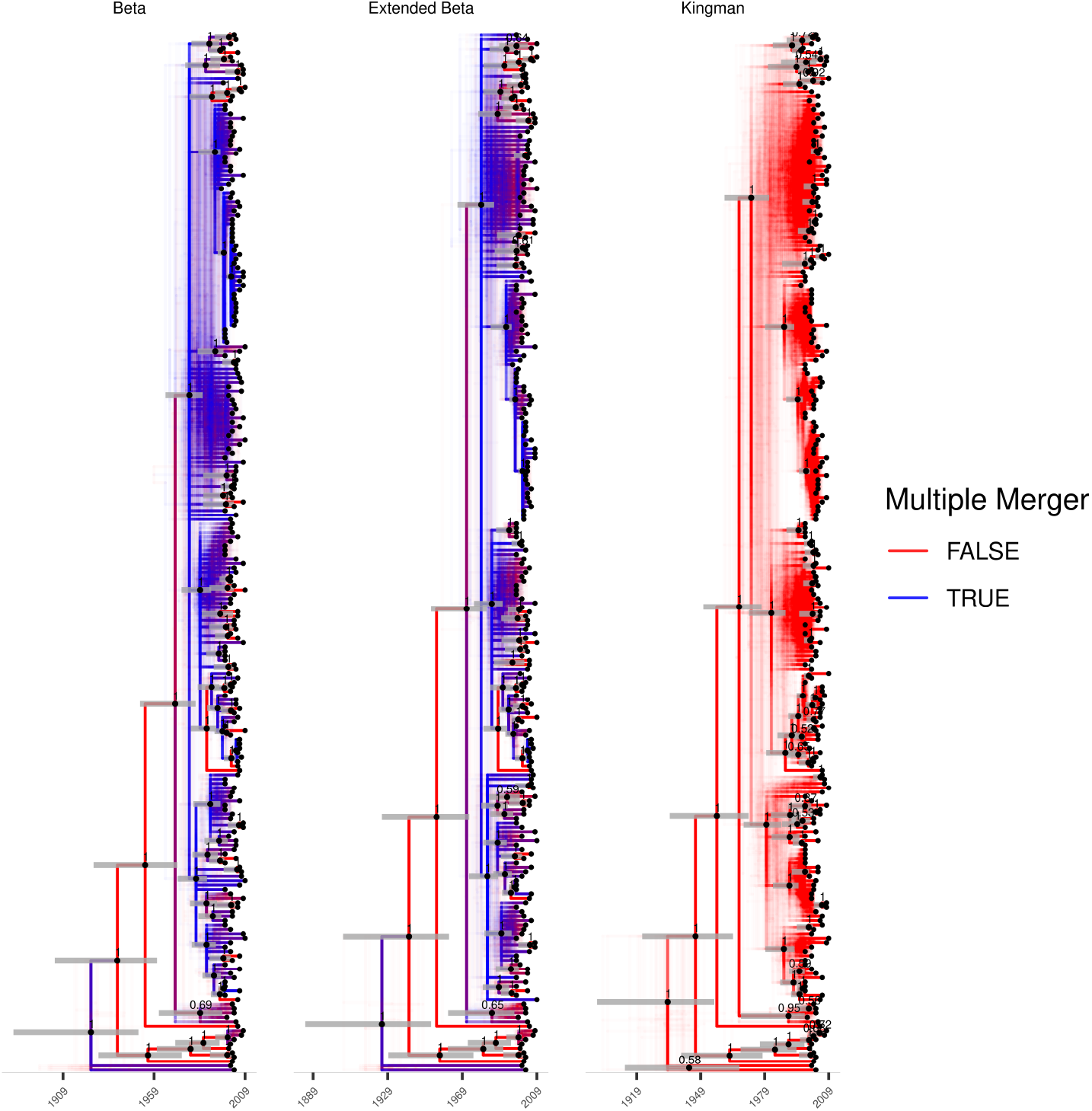
A comparison of three qualitative summaries of posteriors inferred for the the underlying genealogies for the *Eldhom2015* dataset when timed under different coalescent priors. Note that the tip ordering is not exactly the same between the different summaries.

## DISCUSSION

We have presented an approach to reconstructing dated phylogenies with multiple mergers under Lambda-coalescent models. To our knowledge this is the first such approach that scales to real world phylogeny sizes, explicitly reconstructs the underlying multiple merger genealogy, and does not rely on likelihood-free approaches such as Approximate Bayesian Computation. Our focus has been on the implementation of the methodology presented, extensive benchmarks, and applications to real world examples from pathogen phylogenetics. On the other hand we have not yet addressed how this reconstruction can be used to further study of pathogen population genetics. Some implications are relatively straightforward. For example our method could be used as a starting point to do multifurcating skyline plot analysis (Ho and Shapiro 2011). We also envisage that it will lead to other forms of studies becoming possible, for example using genomic data to learn about superspreading, outbreaks, and impacts of selection.

Other Lambda-coalescents models that the ones we used have been studied or derived. While this study was only focused on the Beta-coalescent and its mixture with a Kingman’s coalescent, it is worthwhile to mention some alternatives. In principle there is no reason why the presented methodology would not be applicable to them. A first alternative class of Lambda-coalescent is the extinction-recolonisation Dirac coalescent of (Eldon and Wakeley 2006). In the corresponding forwards in time model all large reproductive events replace a fixed proportion of the population. Each time a potential multiple merger event occurs, a biased coin with probability of heads *p ∈* (0, 1] gets flipped for every extant lineage in the process. Lineages whose coin shows heads all merge into a common ancestor. This model is noteworthy primarily because it was amongst the first to be derived, is simple, and well studied, but it may not be the most biologically plausible.

Another alternative class of Lambda-coalescent is the Durrett-Schweinsberg (DS) coalescent (Durrett and Schweinsberg 2005). This model describes populations undergoing successive hard selective sweeps throughout the genome, and in particular, the hitchhiking effect of those sweeps on a fixed, neutral site. The sweeps are modelled as points in a Poisson process of fixed rate. During a sweep, some ancestral lineages carrying the neutral site of interest can escape the sweep by recombining, while those lineages which don’t recombine will merge to a common ancestor which initiated the sweep. Between individual sweeps the population follows neutral Moran type dynamics. This class of model has recently been found to describe the genetic diversity in cod populations (A–rnason et al. 2023). In general, the measure Λ associated with this class of coalescents takes the form of Λ = *δ*_0_ + Λ_0_ where *δ*_0_ is an atom at 0 responsible for Kingman-like mergers between sweeps and Λ_0_ is a finite measure on [0, 1] without an atom at 0, which drives multiple mergers due to selective sweeps. This model is not directly applicable to pathogen populations, primarily due to traditional recombination being less frequent in bacterial and viral pathogens. An adaptation of this model to bacterial pathogens may be possible but is outside of the scope of this study.

Future studies are needed to investigate what type of Lambda-coalescents best describe pathogen dynamics. The approach presented here is an approximate, albeit explicit approach to Bayesian inference of multiple merger genealogies. It has limitations: for instance, it is not appropriate for studying genealogies spanning geological timescales, as was demonstrated by worsening bias as the number of substitutions per site becomes high (Figure 5). The possibility of extending the inference under Lambda-coalescents to the fully Bayesian setting, incorporating uncertainty about the phylogeny and relying on the phylogenetic likelihood, remains an open problem. Extending some aspects of the parametrisation and construction of multiple merger genealogies presented in this work to the aforementioned setting is straightforward. However, we anticipate that the parametrisation presented here may not be computationally efficient when used in such a setting.

Finally there is the question of extending the approach presented here to, for example, joint estimation of varying effective population size (Ho and Shapiro 2011). This is relevant both for the sake of the past effective population size being an interesting quantity, as well as being relevant from the perspective of statistical robustness. As Menardo et al. (2021) noted, population expansion can be misidentified as a Lambda-coalescent if population growth is not properly accounted for. In order to do this there are separate questions that have to be answered. Firstly, the impact of varying effective population size on the genealogy depends on the forwards-in-time model. Therefore it is first necessary to decide which Lambda-coalescent is the correct one to use for a given scenario. Secondly, adding a non-parametric model for the effective population size will increase the complexity of the inference problem. More efficient MCMC schemes, or other inference tools, would therefore need to be investigated. This might include non-reversible samplers with boundary conditions such as (Bierkens et al. 2023) or Hamiltonian Monte-Carlo methods (Dinh et al. 2017). Note that some phenomena might be indistinguishable from multiple mergers, such as population structure. For example a sufficiently fast expansion of a subpopulation that shares identity by descent (**?**) will likely lead to multiple mergers in the genealogy.

## ACKNOWLEDGEMENTS

DH was supported by the UK Engineering and Physical Sciences Research Council (EPSRC) grant EP/S022244/1 for the EPSRC Centre for Doctoral Training in Mathematics for Real-World Systems

II. JK was supported by EPSRC research grant EP/V049208/1. XD acknowledges funding from the National Institute for Health and Care Research (NIHR) Health Protection Research Unit in Genomics and Enabling Data (NIHR200892).

## References

Bainomugisa, A., E. Lavu, S. Hiashiri, S. Majumdar, A. Honjepari, R. Moke, P. Dakulala, G. A. Hill-Cawthorne, S. Pandey, B. J. Marais, et al. 2018. Multi-clonal evolution of multi-drug-resistant/extensively drug-resistant Mycobacterium tuberculosis in a high-prevalence setting of Papua New Guinea for over three decades. Microbial genomics 4.

Berestycki, N. 2009. Recent progress in coalescent theory.

Bertoin, J. and J.-F. Le Gall. 2000. The Bolthausen–Sznitman coalescent and the genealogy of continuous-state branching processes. Probability Theory and Related Fields 117:249–266.

Biek, R., O. G. Pybus, J. O. Lloyd-Smith, and X. Didelot. 2015. Measurably Evolving Pathogens in the Genomic Era. Trends in Ecology & Evolution 30:306–313 publisher: Elsevier Ltd.

Bierkens, J., S. Grazzi, F. v. d. Meulen, and M. Schauer. 2023. Sticky PDMP samplers for sparse and local inference problems. Statistics and Computing 33:8.

Billera, L. J., S. P. Holmes, and K. Vogtmann. 2001. Geometry of the Space of Phylogenetic Trees. Advances in Applied Mathematics 27:733–767.

Birkner, M., J. Blath, and B. Eldon. 2013. Statistical Properties of the Site-Frequency Spectrum Associated with Λ-Coalescents. Genetics 195:1037–1053.

Bouckaert, R. R. 2010. DensiTree: making sense of sets of phylogenetic trees. Bioinformatics 26:1372–1373.

Bromham, L. and D. Penny. 2003. The modern molecular clock. Nature Reviews Genetics 4:216–224.

Charlesworth, B. 2009. Fundamental concepts in genetics: Effective population size and patterns of molecular evolution and variation. Nature Reviews Genetics 10:195–205.

Cvijović, I., B. H. Good, and M. M. Desai. 2018. The Effect of Strong Purifying Selection on Genetic Diversity. Genetics 209:1235–1278.

Desai, M. M., A. M. Walczak, and D. S. Fisher. 2013. Genetic Diversity and the Structure of Genealogies in Rapidly Adapting Populations. Genetics 193:565–585.

Didelot, X., N. J. Croucher, S. D. Bentley, S. R. Harris, and D. J. Wilson. 2018. Bayesian inference of ancestral dates on bacterial phylogenetic trees. Nucleic Acids Res. 46:e134–e134.

Didelot, X., B. Pang, Z. Zhou, A. McCann, P. Ni, D. Li, M. Achtman, and B. Kan. 2015. The Role of China in the Global Spread of the Current Cholera Pandemic. PLoS Genetics 11:e1005072.

Didelot, X. and J. Parkhill. 2022. A Scalable Analytical Approach from Bacterial Genomes to Epidemiology. Philosophical Transactions of the Royal Society B: Biological Sciences 377:20210246 publisher: Cold Spring Harbor Laboratory.

Didelot, X., I. Siveroni, and E. M. Volz. 2021. Additive uncorrelated relaxed clock models for the dating of genomic epidemiology phylogenies. Mol. Biol. Evol. 38:307–317.

Dinh, V., A. Bilge, C. Zhang, and F. A. M. Iv. 2017. Probabilistic Path Hamiltonian Monte Carlo. Pages 1009–1018 in Proceedings of the 34th International Conference on Machine Learning PMLR iSSN: 2640-3498.

Donnelly, P. and T. G. Kurtz. 1999. Particle Representations for Measure-Valued Population Models. The Annals of Probability 27:166–205 publisher: Institute of Mathematical Statistics.

Dorman, M. J., D. Domman, T. Poklepovich, C. Tolley, G. Zolezzi, L. Kane, M. R. Vinñas, M. Panagópulo, M. Moroni, N. Binsztein, M. I. Caffer, S. Clare, G. Dougan, G. P. C. Salmond, J. Parkhill, J. Campos, and N. R. Thomson. 2020. Genomics of the Argentinian cholera epidemic elucidate the contrasting dynamics of epidemic and endemic Vibrio cholerae. Nature Communications 11:4918.

Drummond, A. J., G. K. Nicholls, A. G. Rodrigo, and W. Solomon. 2002. Estimating mutation parameters, population history and genealogy simultaneously from temporally spaced sequence data. Genetics 161:1307–1320.

Drummond, A. J., O. G. Pybus, A. Rambaut, R. Forsberg, and A. G. Rodrigo. 2003. Measurably Evolving Populations. Trends in Ecology and Evolution 18:481–488.

Durrett, R. and J. Schweinsberg. 2005. A coalescent model for the effect of advantageous mutations on the genealogy of a population. Stochastic Processes and their Applications 115:1628–1657.

Eldholm, V., J. Monteserin, A. Rieux, B. Lopez, B. Sobkowiak, V. Ritacco, and F. Balloux. 2015. Four decades of transmission of a multidrug-resistant Mycobacterium tuberculosis outbreak strain. Nat. Commun. 6:7119 publisher: Nature Publishing Group.

Eldon, B. and W. Stephan. 2023. Sweepstakes reproduction facilitates rapid adaptation in highly fecund populations. Molecular Ecology n/a eprint: https://onlinelibrary.wiley.com/doi/pdf/10.1111/mec.16903.

Eldon, B. and J. Wakeley. 2006. Coalescent Processes When the Distribution of Offspring Number Among Individuals Is Highly Skewed. Genetics 172:2621–2633.

Gabry, J., D. Simpson, A. Vehtari, M. Betancourt, and A. Gelman. 2019. Visualization in Bayesian workflow. Journal of the Royal Statistical Society Series A: Statistics in Society 182:389–402.

George, E. I. and R. E. McCulloch. 1993. Variable Selection via Gibbs Sampling. Journal of the American Statistical Association 88:881–889 publisher: Taylor & Francis eprint: https://www.tandfonline.com/doi/pdf/10.1080/01621459.1993.10476353.

Green, P. J. 1995. Reversible Jump Markov Chain Monte Carlo Computation and Bayesian Model Determination. Biometrika 82:711–732.

Guindon, S., J.-F. Dufayard, V. Lefort, M. Anisimova, W. Hordijk, and O. Gascuel. 2010. New Algorithms and Methods to Estimate Maximum-Likelihood Phylogenies: Assessing the Performance of PhyML 3.0. Systematic biology 59:307–21.

Gómez-Carballa, A., J. Pardo-Seco, X. Bello, F. Martinón-Torres, and A. Salas. 2021. Superspreading in the emergence of COVID-19 variants. Trends in Genetics 37:1069–1080.

Hasegawa, M., H. Kishino, and T. Yano. 1985. Dating of the Human-Ape Splitting by a Molecular Clock of Mitochondrial DNA. Journal of molecular evolution 22:160–174.

Helekal, D., A. Ledda, E. Volz, D. Wyllie, and X. Didelot. 2021. Bayesian Inference of Clonal Expansions in a Dated Phylogeny. Systematic Biology Page syab095.

Ho, S. Y. W. and B. Shapiro. 2011. Skyline-plot methods for estimating demographic history from nucleotide sequences. Mol. Ecol. Resour. 11:423–434 iSBN: 1755-0998 (Electronic)\backslashr1755-098X (Linking).

Hoscheit, P. and O. G. Pybus. 2019. The multifurcating skyline plot. Virus Evol. 5:1–10 iSBN: 000000080013.

Ji, X., A. A. Fisher, S. Su, J. L. Thorne, B. Potter, P. Lemey, G. Baele, and M. A. Suchard. 2021. Scalable Bayesian divergence time estimation with ratio transformations. ArXiv:2110.13298 [q-bio, stat].

Kingman, J. F. C. 1982. The coalescent. Stochastic Processes and their Applications 13:235–248.

Kuhn, T. S., A. O. Mooers, and G. H. Thomas. 2011. A simple polytomy resolver for dated phylogenies. Methods in Ecology and Evolution 2:427–436 eprint: https://onlinelibrary.wiley.com/doi/pdf/10.1111/j.2041-210X.2011.00103.x.

Kukla, J. and M. Möhle. 2018. On the block counting process and the fixation line of the Bolthausen– Sznitman coalescent. Stochastic Processes and their Applications 128:939–962.

Lee, R. S., N. Radomski, J.-F. Proulx, I. Levade, B. J. Shapiro, F. McIntosh, H. Soualhine, D. Menzies, and M. A. Behr. 2015. Population genomics of Mycobacterium tuberculosis in the Inuit. Proceedings of the National Academy of Sciences 112:13609–13614.

Lemieux, J. E., K. J. Siddle, B. M. Shaw, C. Loreth, S. F. Schaffner, A. Gladden-Young, G. Adams, T. Fink, C. H. Tomkins-Tinch, L. A. Krasilnikova, K. C. DeRuff, M. Rudy, M. R. Bauer, K. A. Lagerborg, E. Normandin, S. B. Chapman, S. K. Reilly, M. N. Anahtar, A. E. Lin, A. Carter, C. Myhrvold, M. E. Kemball, S. Chaluvadi, C. Cusick, K. Flowers, A. Neumann, F. Cerrato, M. Farhat, D. Slater, J. B. Harris, J. Branda, D. Hooper, J. M. Gaeta, T. P. Baggett, J. O’Connell, A. Gnirke, T. D. Lieberman, A. Philippakis, M. Burns, C. M. Brown, J. Luban, E. T. Ryan, S. E. Turbett, R. C. LaRocque, W. P. Hanage, G. R. Gallagher, L. C. Madoff, S. Smole, V. M. Pierce, E. Rosenberg, P. C. Sabeti, D. J. Park, and B. L. Maclnnis. 2020. Phylogenetic analysis of SARS-CoV-2 in the Boston area highlights the role of recurrent importation and superspreading events. preprint Epidemiology.

Lewis, P. O., M. T. Holder, and K. E. Holsinger. 2005. Polytomies and Bayesian Phylogenetic Inference. Systematic Biology 54:241–253.

Lin, G. N., C. Zhang, and D. Xu. 2011. Polytomy identification in microbial phylogenetic reconstruction. BMC Systems Biology 5:S2.

Matuszewski, S., M. E. Hildebrandt, G. Achaz, and J. D. Jensen. 2018. Coalescent Processes with Skewed Offspring Distributions and Nonequilibrium Demography. Genetics 208:323–338.

Menardo, F., S. Gagneux, and F. Freund. 2021. Multiple Merger Genealogies in Outbreaks of Mycobacterium tuberculosis. Molecular Biology and Evolution 38:290–306.

Metropolis, N., A. W. Rosenbluth, M. N. Rosenbluth, A. H. Teller, and E. Teller. 1953. Equation of State Calculations by Fast Computing Machines. Journal of Chemical Physics 21:1087–1092 aDS Bibcode: 1953JChPh..21.1087M.

Minh, B. Q., H. A. Schmidt, O. Chernomor, D. Schrempf, M. D. Woodhams, A. von Haeseler, and R. Lanfear. 2020. IQ-TREE 2: New Models and Efficient Methods for Phylogenetic Inference in the Genomic Era. Molecular Biology and Evolution 37:1530–1534.

Mutreja, A., D. W. Kim, N. R. Thomson, T. R. Connor, J. H. Lee, S. Kariuki, N. J. Croucher, S. Y. Choi, S. R. Harris, M. Lebens, S. K. Niyogi, E. J. Kim, T. Ramamurthy, J. Chun, J. L. N. Wood, J. D. Clemens, C. Czerkinsky, G. B. Nair, J. Holmgren, J. Parkhill, and G. Dougan. 2011. Evidence for several waves of global transmission in the seventh cholera pandemic. Nature 477:462–465.

Neher, R. A. and O. Hallatschek. 2013. Genealogies of rapidly adapting populations. Proceedings of the National Academy of Sciences 110:437–442 publisher: Proceedings of the National Academy of Sciences.

Paradis, E. and K. Schliep. 2019. ape 5.0: an environment for modern phylogenetics and evolutionary analyses in R. Bioinformatics 35:526–528 publisher: Oxford University Press.

Pitman, J. 1999. Coalescents With Multiple Collisions. The Annals of Probability 27:1870–1902 publisher: Institute of Mathematical Statistics.

Rambaut, A. and N. C. Grass. 1997. Seq-Gen: an application for the Monte Carlo simulation of DNA sequence evolution along phylogenetic trees. Bioinformatics 13:235–238.

Robinson, D. F. and L. R. Foulds. 1981. Comparison of phylogenetic trees. Mathematical Biosciences 53:131–147.

Sagitov, S. 1999. The general coalescent with asynchronous mergers of ancestral lines. Journal of Applied Probability 36:1116–1125 publisher: Cambridge University Press.

Sagulenko, P., V. Puller, and R. A. Neher. 2018. TreeTime: Maximum likelihood phylodynamic analysis. Virus Evol. 4:vex042.

Schweinsberg, J. 2003. Coalescent processes obtained from supercritical Galton–Watson processes. Stochastic Processes and their Applications 106:107–139.

Schweinsberg, J. 2017. Rigorous results for a population model with selection II: genealogy of the population. Electronic Journal of Probability 22:1–54 publisher: Institute of Mathematical Statistics and Bernoulli Society.

Stamatakis, A. 2014. RAxML version 8: a tool for phylogenetic analysis and post-analysis of large phylogenies. Bioinformatics 30:1312–1313.

Tellier, A. and C. Lemaire. 2014. Coalescence 2.0: a multiple branching of recent theoretical developments and their applications. Molecular ecology 23:2637–2652.

To, T.-H., M. Jung, S. Lycett, and O. Gascuel. 2016. Fast Dating Using Least-Squares Criteria and Algorithms. Systematic Biology 65:82–97.

Vehtari, A., A. Gelman, D. Simpson, B. Carpenter, and P. C. Burkner. 2021. Rank-Normalization, Folding, and Localization: An Improved R hat for Assessing Convergence of MCMC. Bayesian Anal. 16:667–718 eprint: arXiv:1903.08008v5.

Volz, E. M. and S. D. W. Frost. 2017. Scalable Relaxed Clock Phylogenetic Dating. Virus Evolution 3:vex025.

Yang, Z. and B. Rannala. 1997. Bayesian Phylogenetic Inference using DNA Sequences: A Markov Chain Monte Carlo Method. Molecular Biology and Evolution 14:717–724 publisher: SMBE.

Yu, G., D. K. Smith, H. Zhu, Y. Guan, and T. T. Y. Lam. 2017. Ggtree: an R Package for Visualization and Annotation of Phylogenetic Trees With Their Covariates and Other Associated Data. Methods Ecol. Evol. 8:28–36.

Árnason, E., J. Koskela, K. Halldórsdóttir, and B. Eldon. 2023. Sweepstakes reproductive success via pervasive and recurrent selective sweeps. eLife 12:e80781 publisher: eLife Sciences Publications, Ltd.

