## Supplementary Material for "Inference of multiple mergers while dating a pathogen phylogeny"

### Inference of multiple mergers while dating a pathogen phylogeny
#### Supplementary Material

David Helekal, Jere Koskela, Xavier Didelot

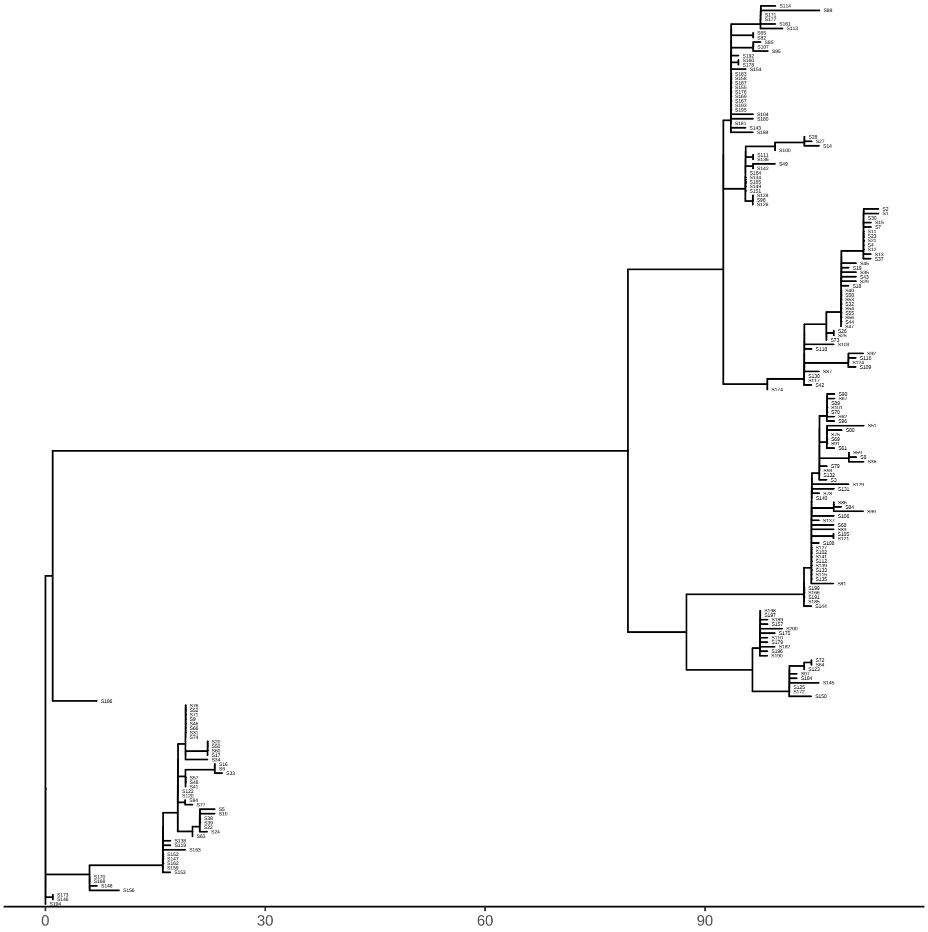


Figure S1: Input phylogeny for the Beta-coalescent example.

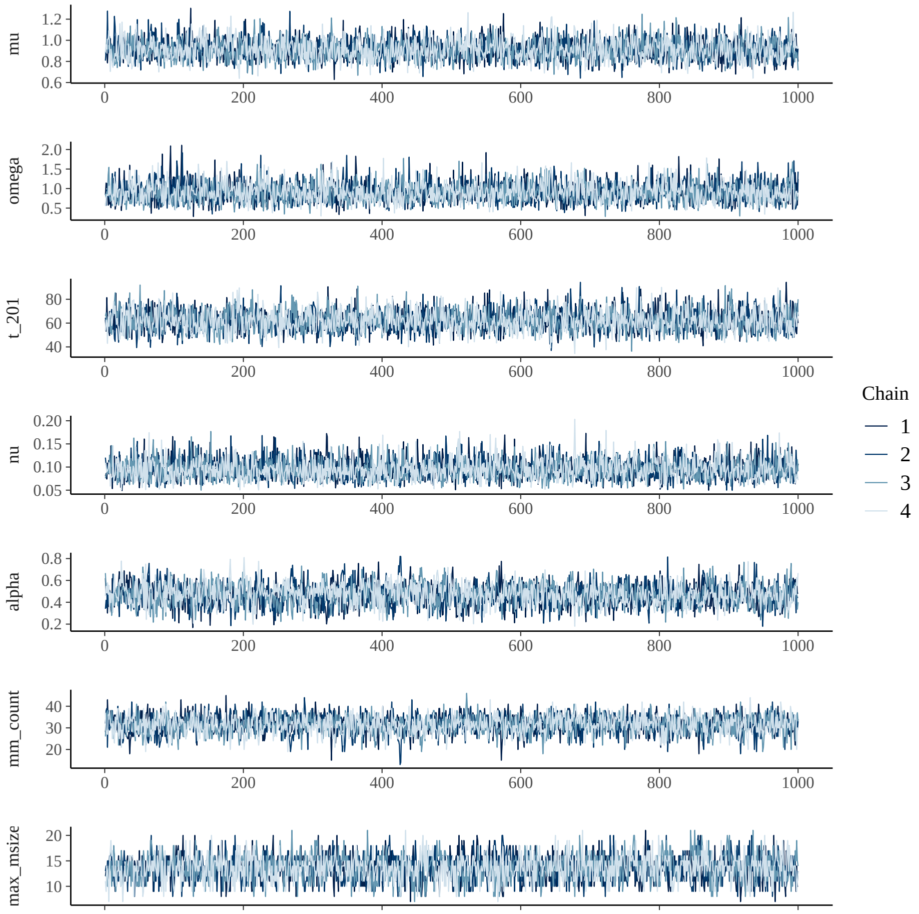


Figure S2: Parameter and diagnostic quantity traces for the Beta-coalescent example.

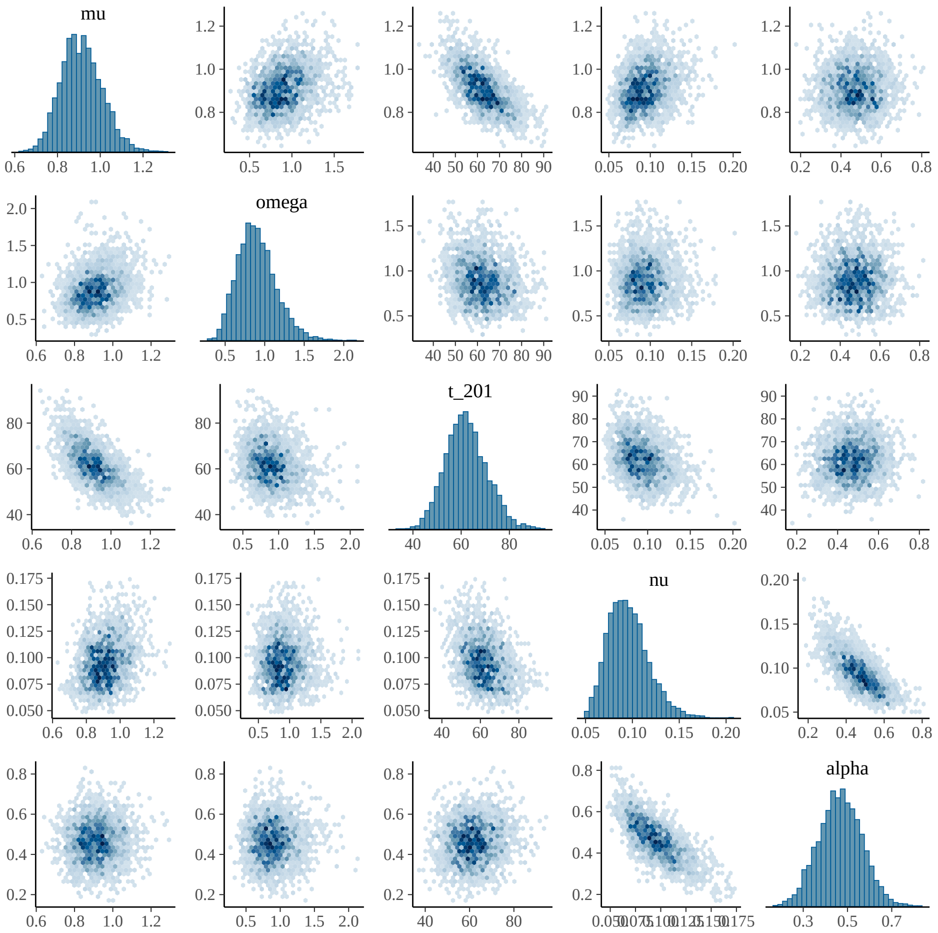


Figure S3: Parameter posterior distributions for the Beta-coalescent example.

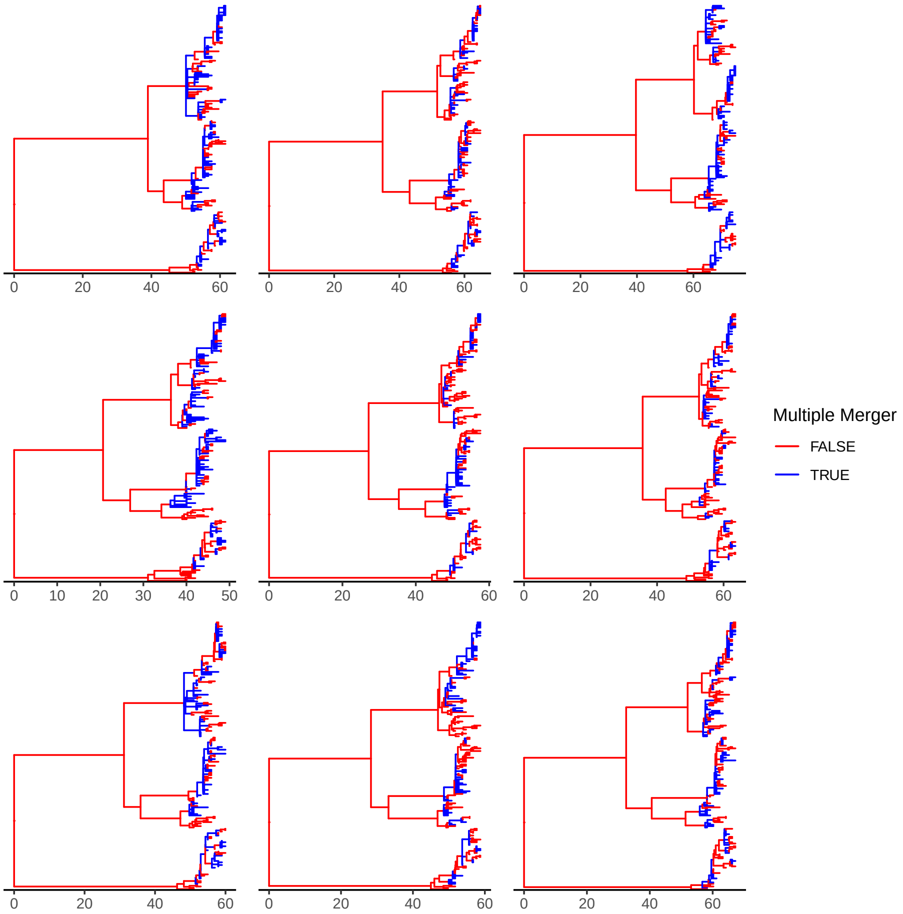


Figure S4: Nine realisations of the underlying genealogy sampled from the posterior estimated for the Beta-coalescent example. Branches are colored according to presence of multiple merger directly above.

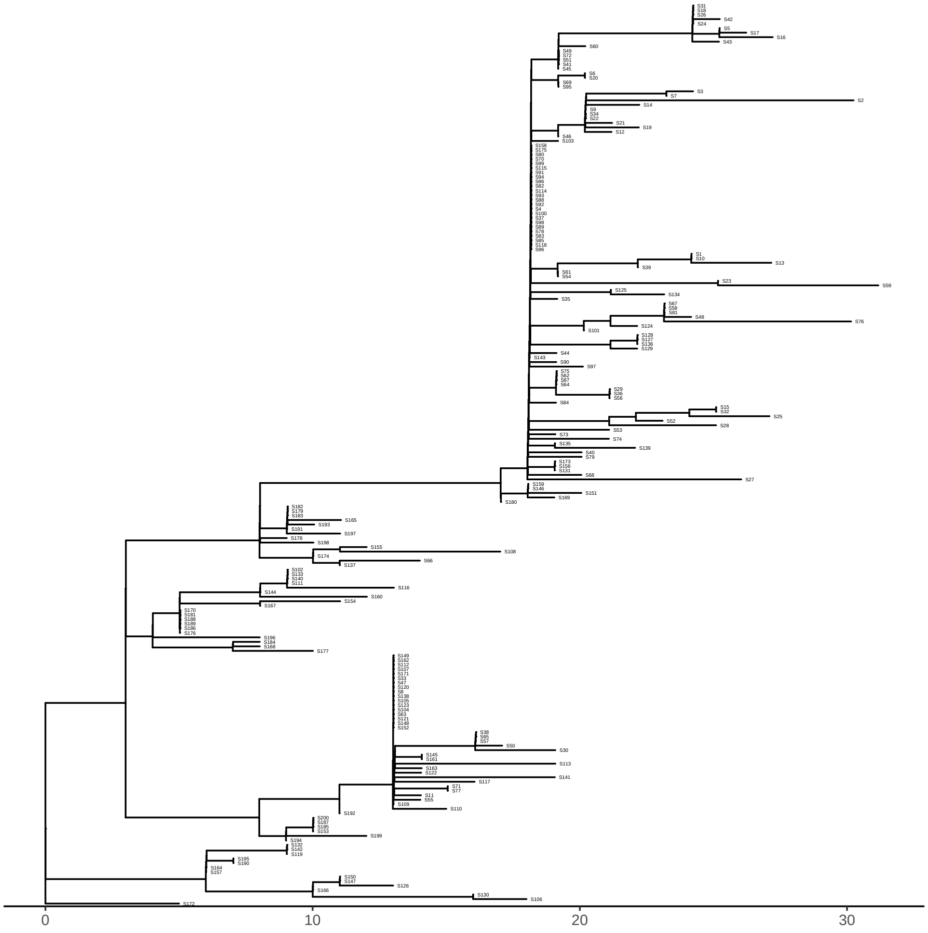


Figure S5: Input phylogeny for the extended Beta-coalescent example.

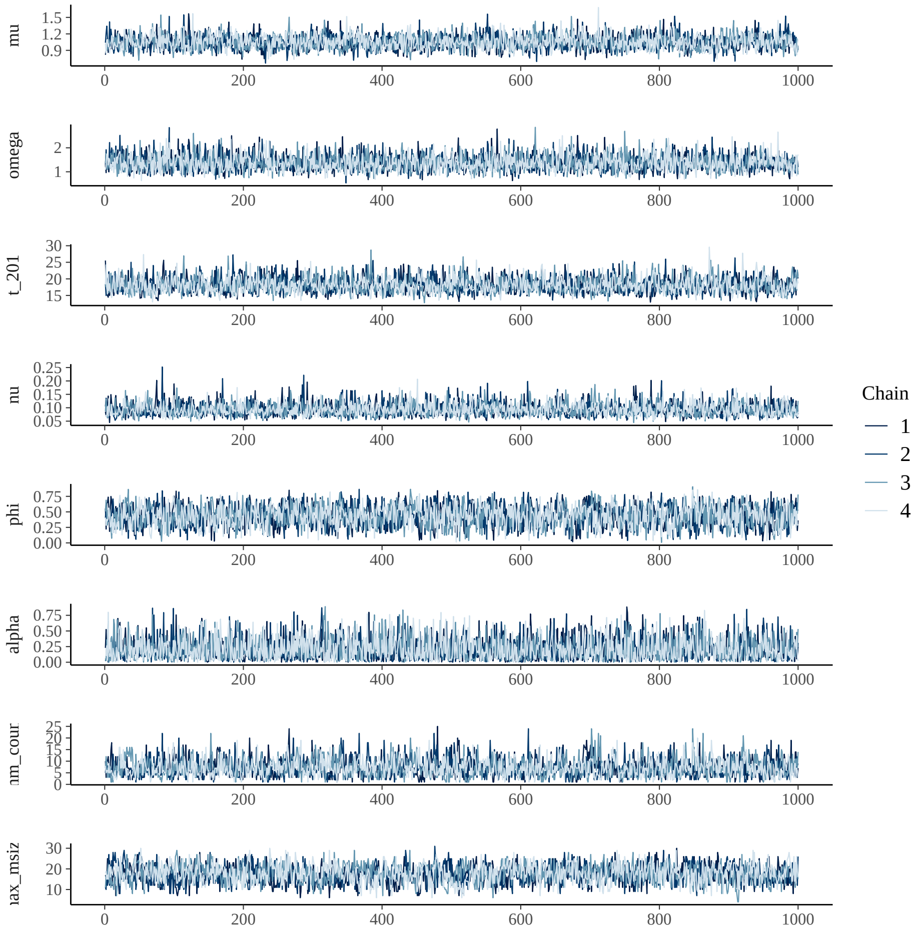


Figure S6: Parameter and diagnostic quantity traces for the extended Beta-coalescent example.

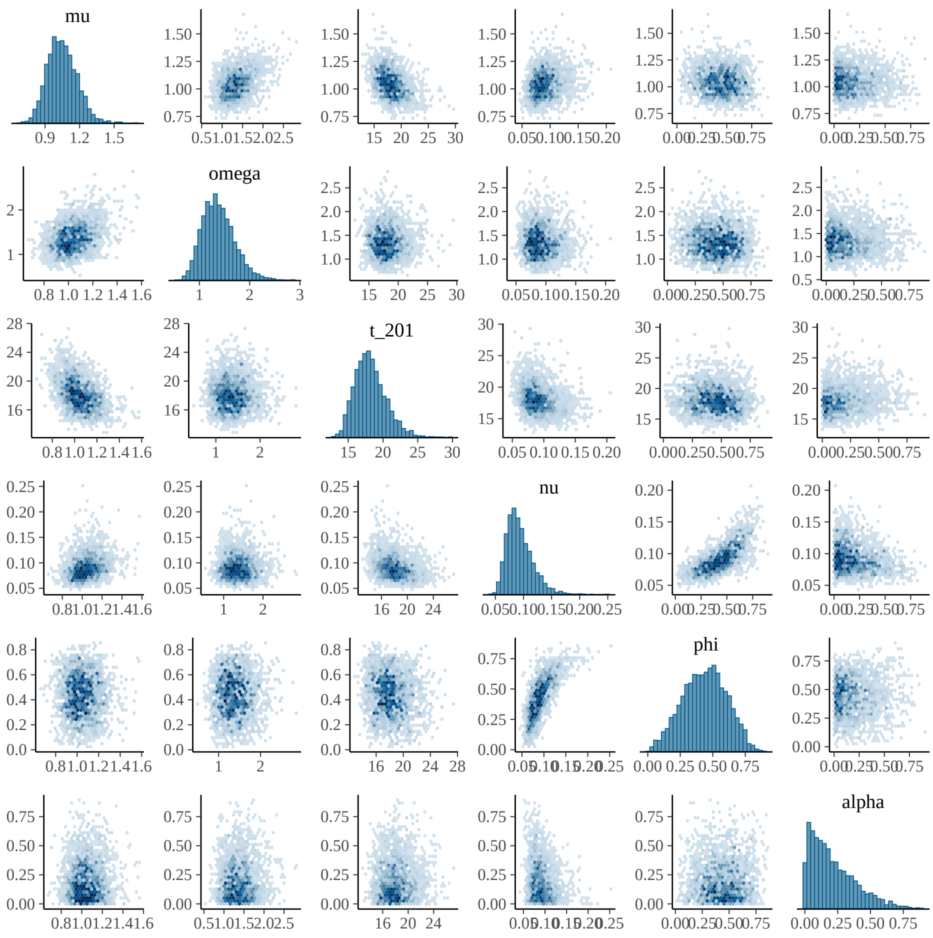


Figure S7: Parameter posterior distributions for the extended Beta-coalescent example.

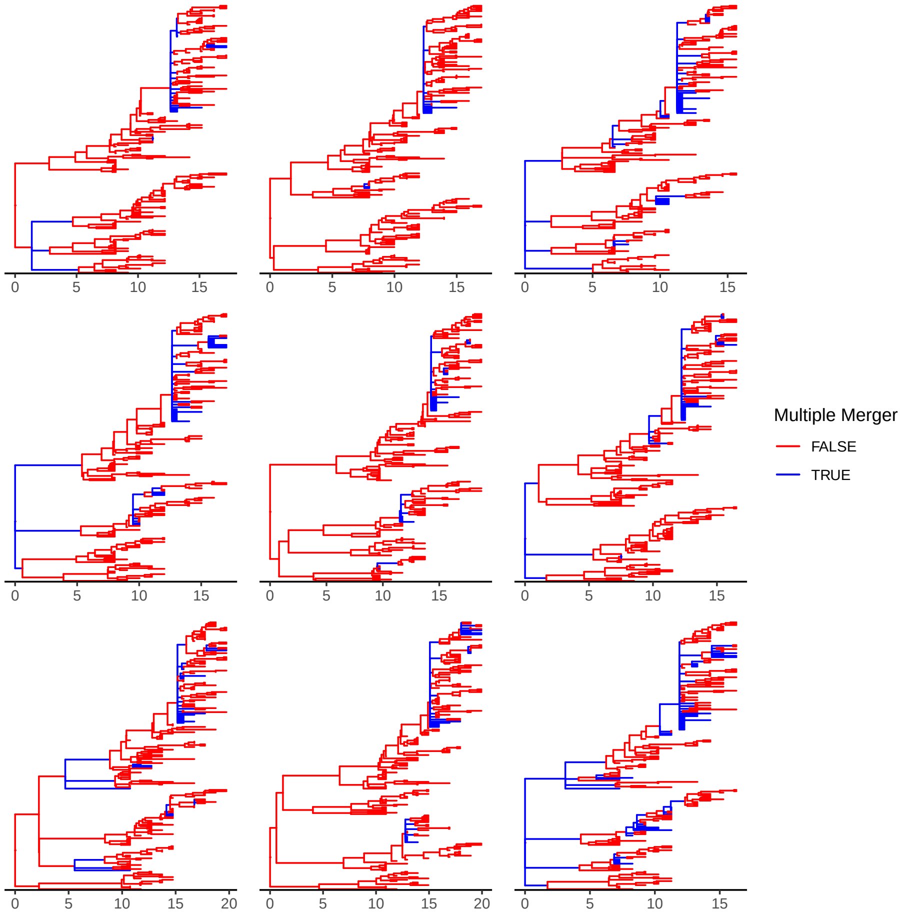


Figure S8: Nine realisations of the underlying genealogy sampled from the posterior estimated for the extended Beta-coalescent example. Branches are colored according to presence of multiple merger directly above.

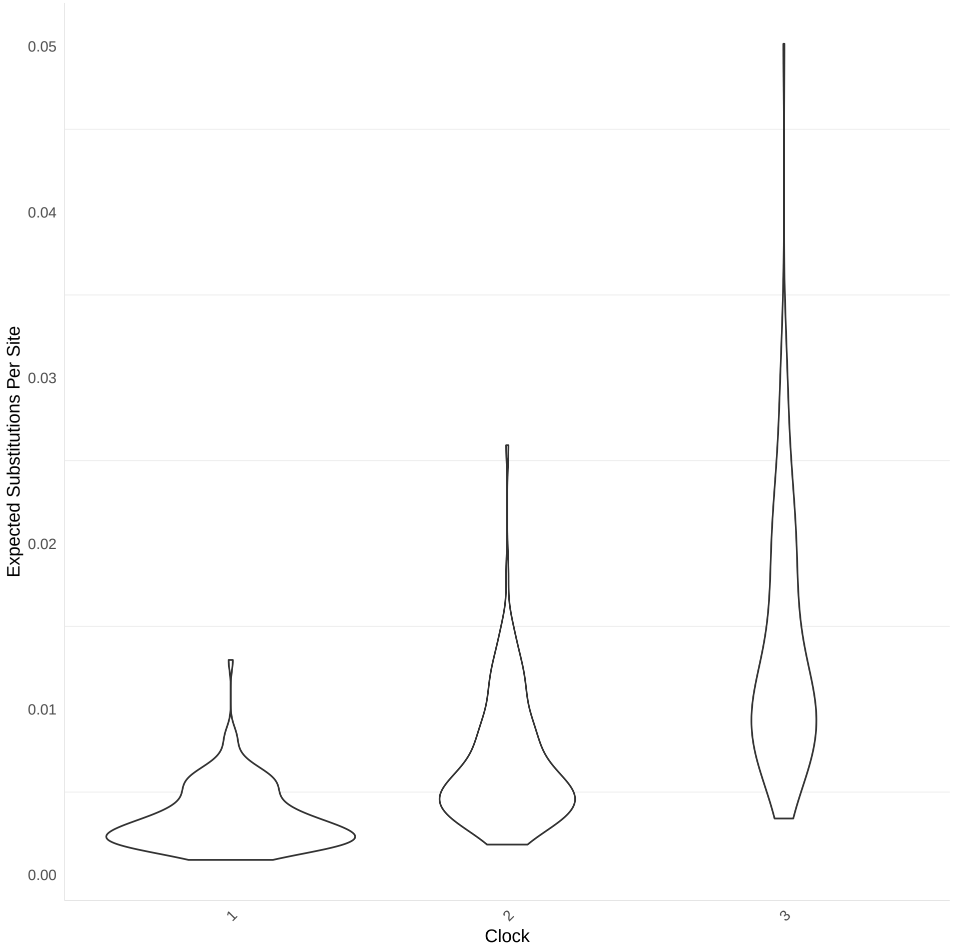


Figure S9: Expected number of substitutions for simulated sequences used for the analysis under the Beta-coalescent.

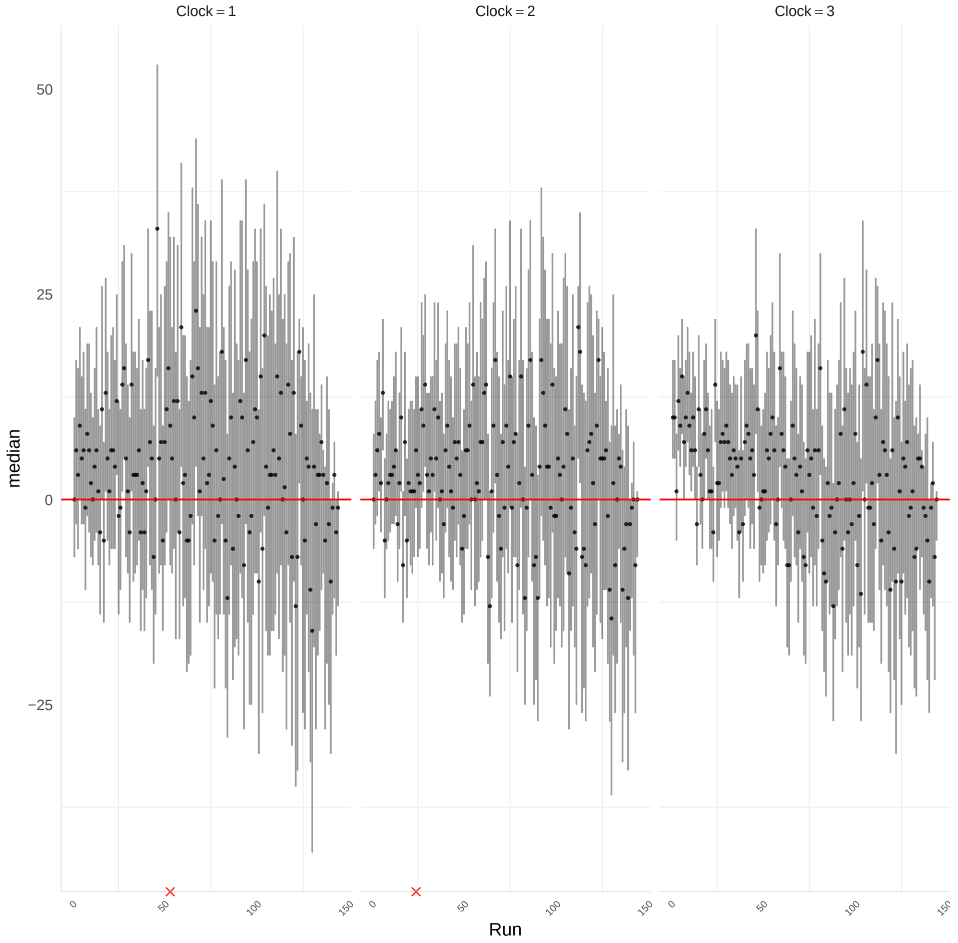


Figure S10: Number of nodes in posterior samples minus the true number of nodes in the simulated genealogy for the analysis under the Beta-coalescent. Red crosses indicate runs that indicated unsatisfactory mixing.

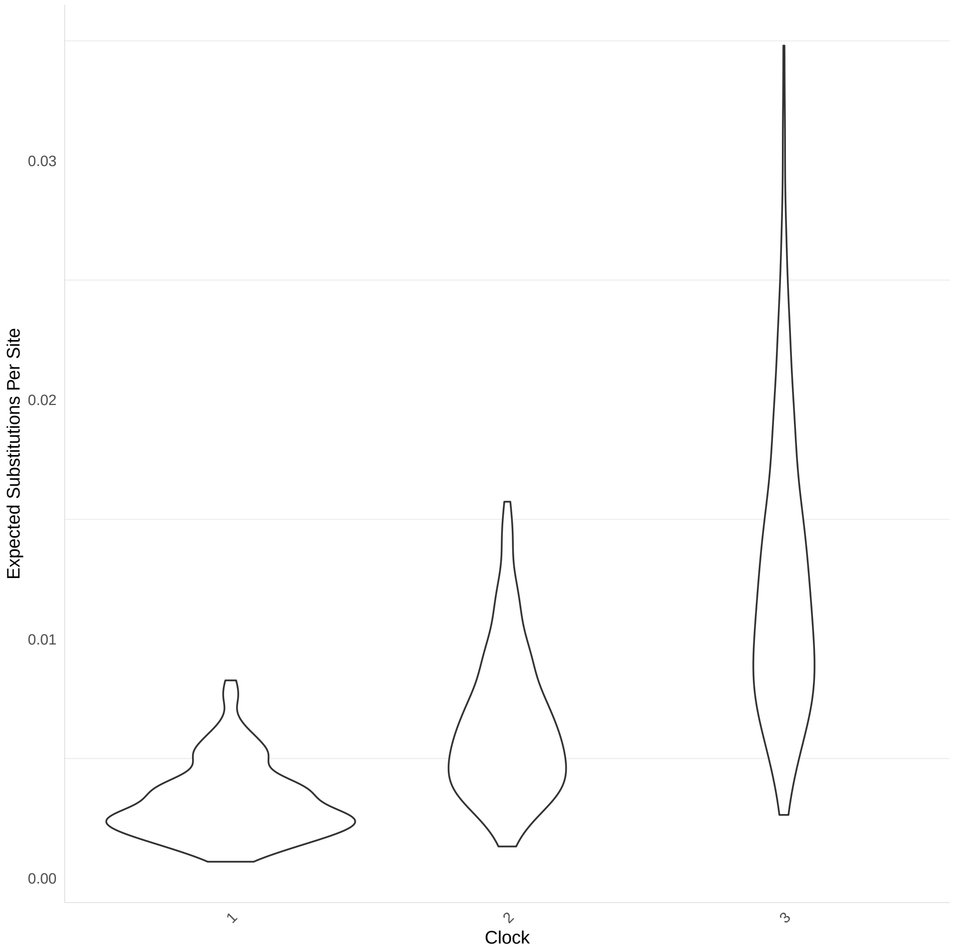


Figure S11: Expected number of substitutions for simulated sequences used for the analysis under the extended Beta-coalescent. Red crosses indicate runs that indicated unsatisfactory mixing.

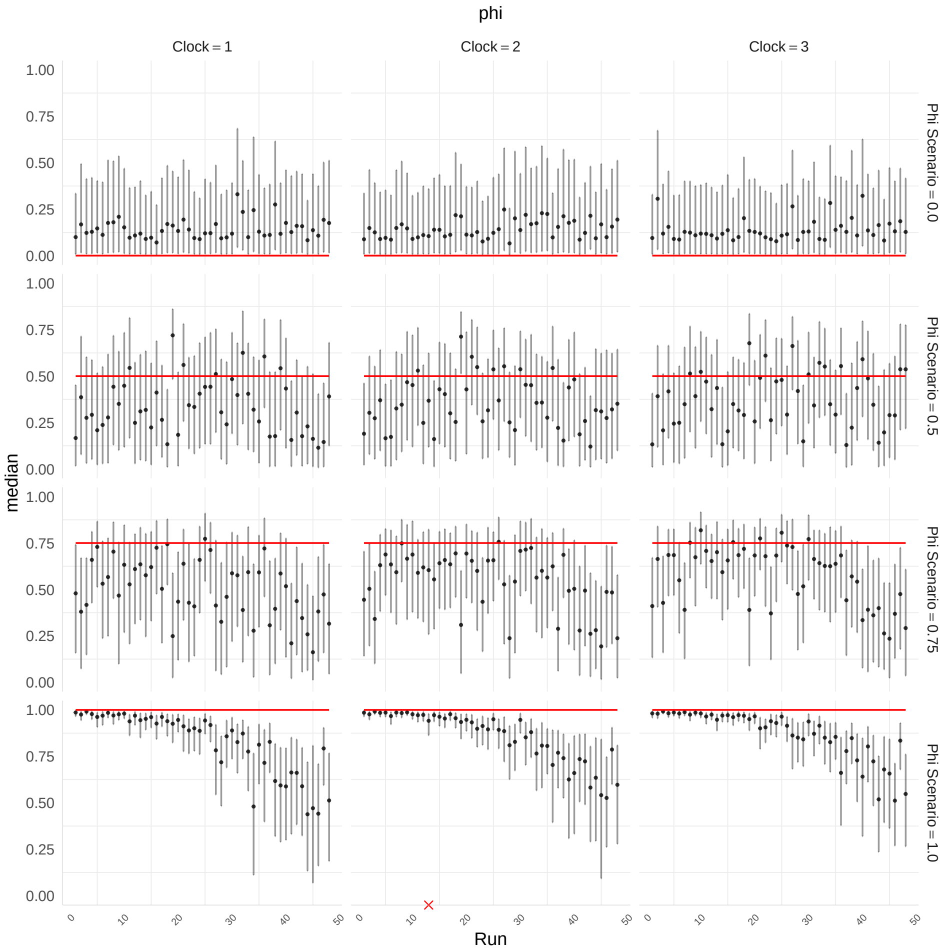


Figure S12: Posterior summaries for the $\phi$ parameter for the analysis under the extended Beta-coalescent. Red crosses indicate runs that indicated unsatisfactory mixing.

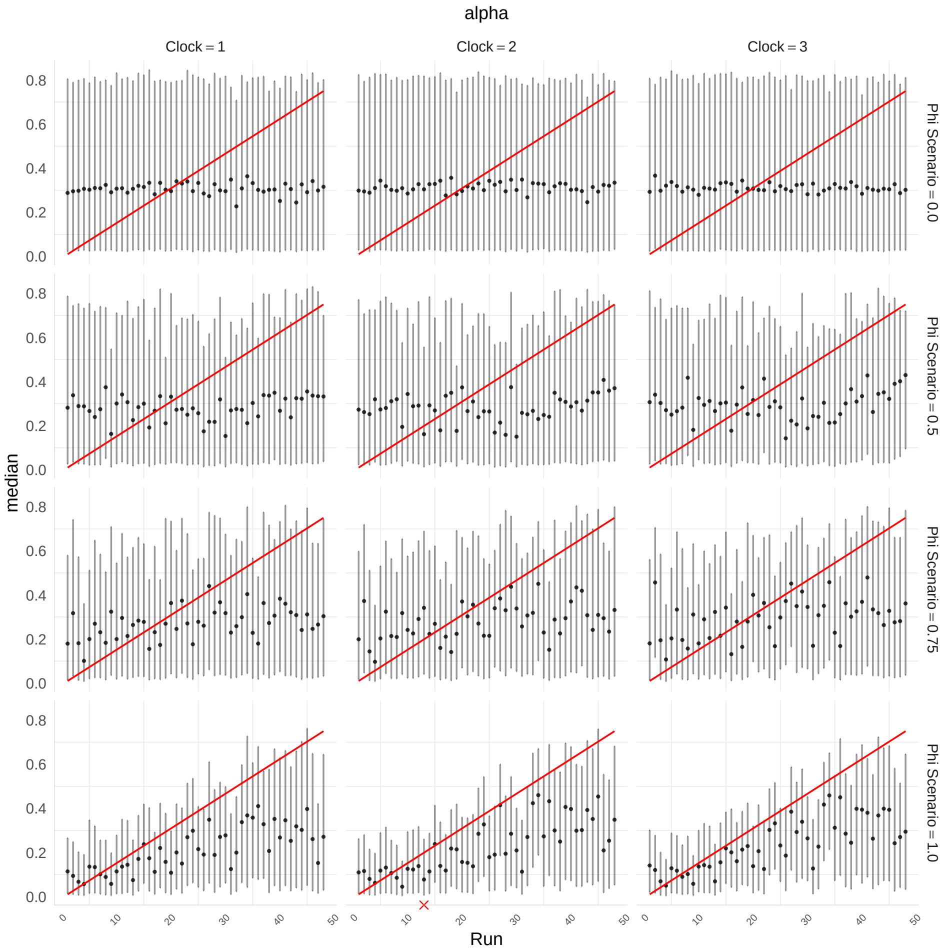


Figure S13: Posterior summaries for the $\alpha^*$ parameter for the analysis under the extended Beta-coalescent.

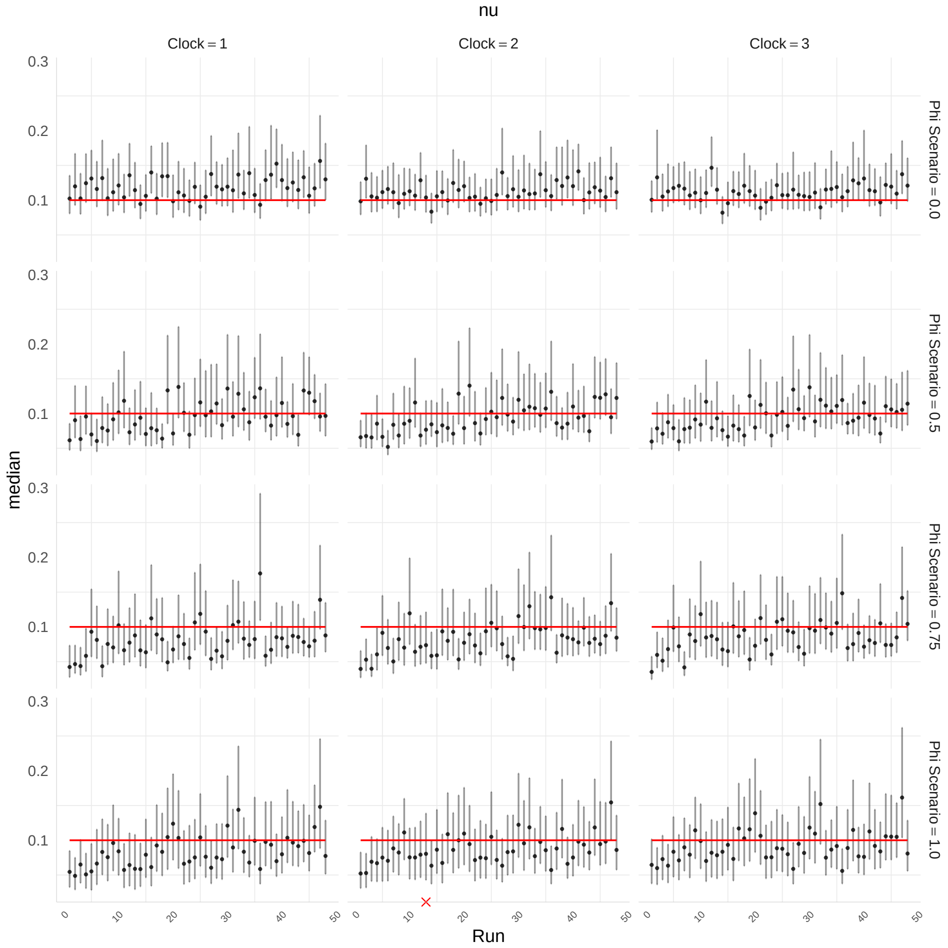


Figure S14: Posterior summaries for the $\nu$ parameter for the analysis under the extended Beta-coalescent. Red crosses indicate runs that indicated unsatisfactory mixing.

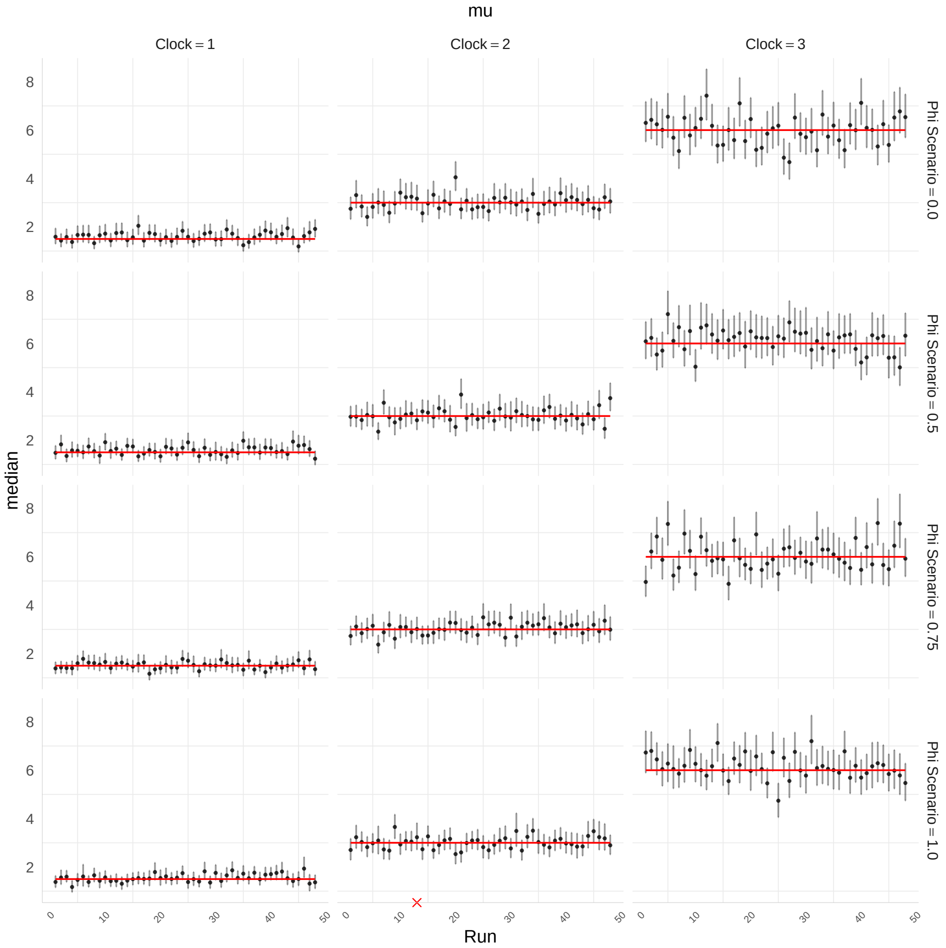


Figure S15: Posterior summaries for the $\mu$ parameter for the analysis under the extended Beta-coalescent. Red bold line indicates simulation value. Red crosses indicate runs that indicated unsatisfactory mixing.

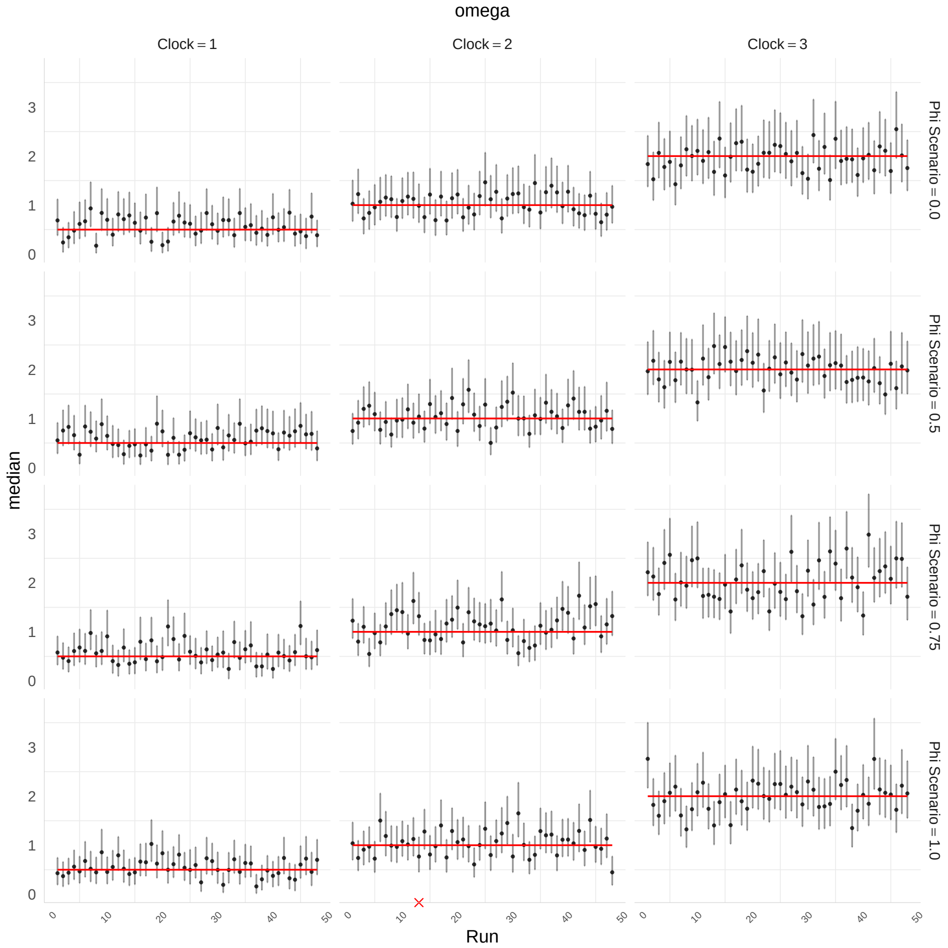


Figure S16: Posterior summaries for the $\omega$ parameter for the analysis under the extended Beta-coalescent. Red bold line indicates simulation value. Red crosses indicate runs that indicated unsatisfactory mixing.

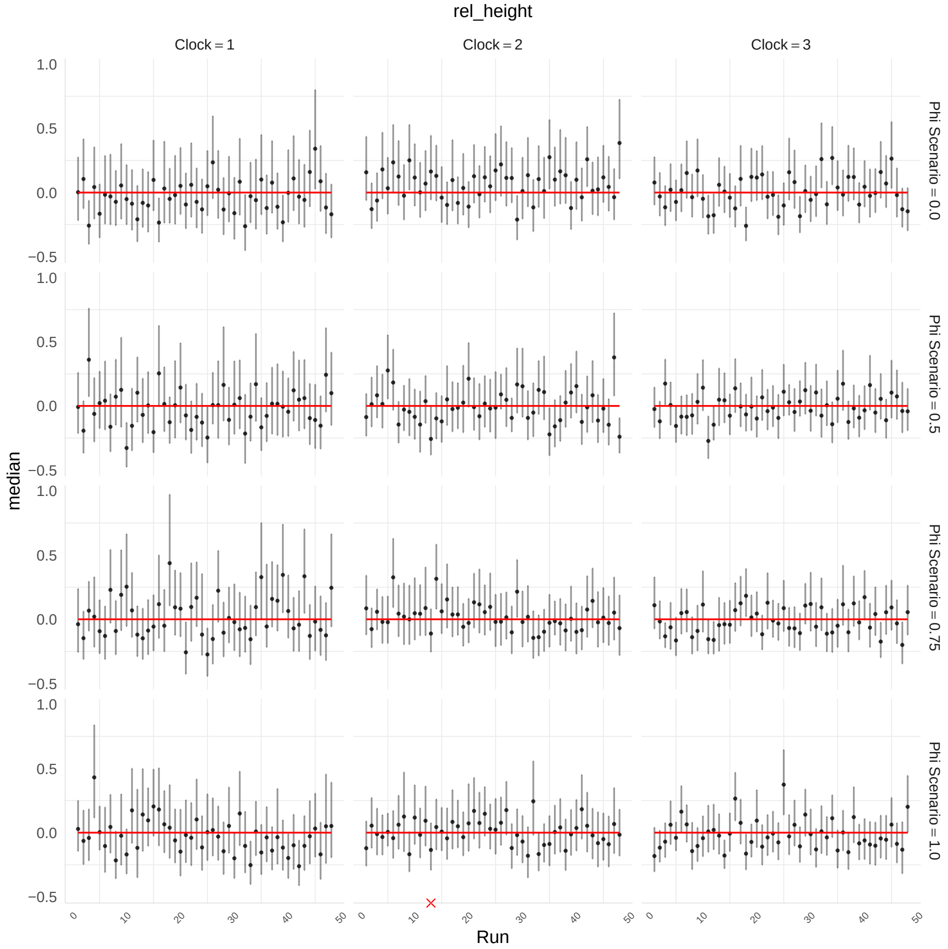


Figure S17: Posterior summary of relative tree heights for the analysis under the extended Beta-coalescent. Red crosses indicate runs that indicated unsatisfactory mixing.

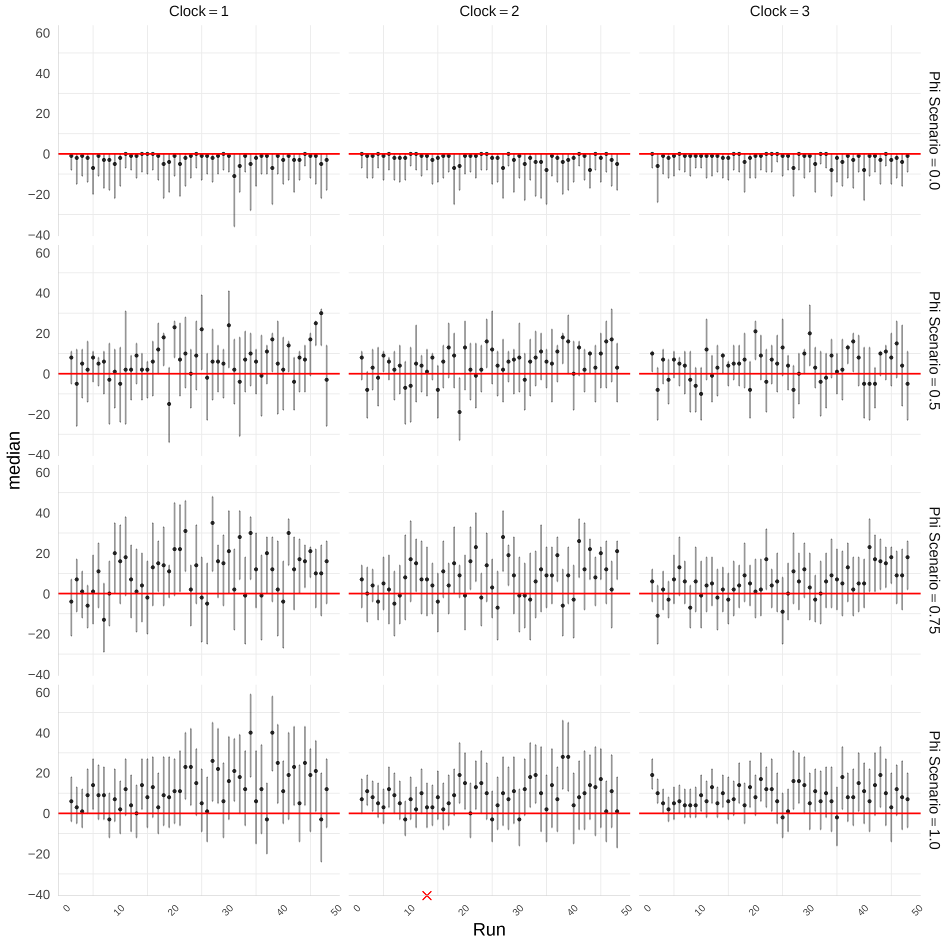


Figure S18: Number of nodes in the posterior samples minus the number of nodes in the simulated genealogy for the analysis under the extended Beta-coalescent. Red crosses indicate runs that indicated unsatisfactory mixing.

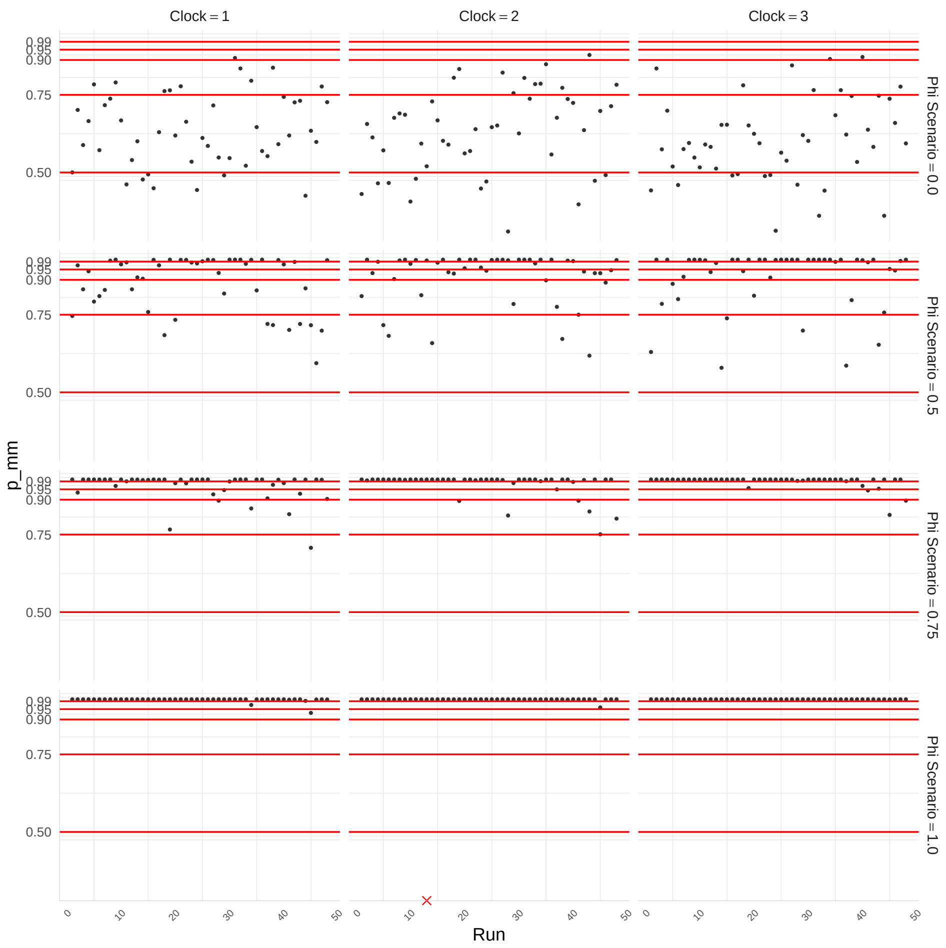


Figure S19: Estimated probabilities that a tree sampled from the posterior contains at least one multi-merger event for the analysis under the extended Beta-coalescent. Red lines indicate the 50%, 75%, 90%, 95%, and 99% thresholds. Red crosses indicate runs that indicated unsatisfactory mixing.

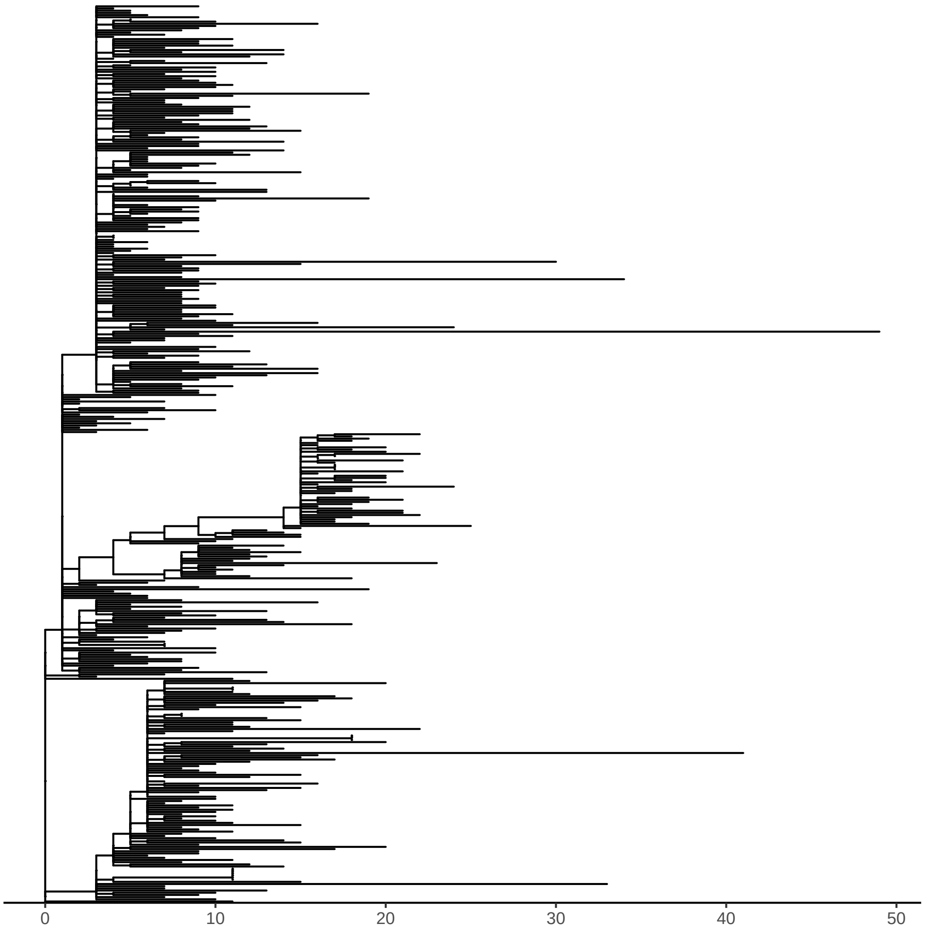


Figure S20: Input phylogeny for the *Vibrio cholerae* case study.

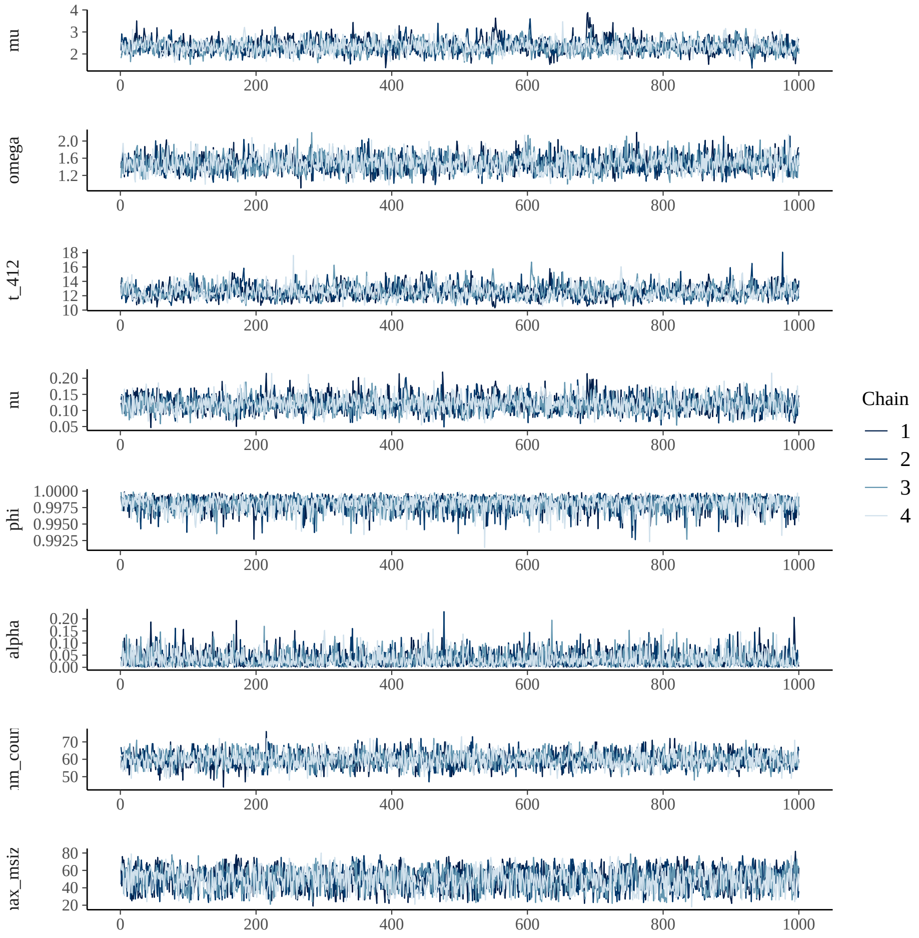


Figure S21: Parameter and diagnostic quantity traces for the analysis of the *Vibrio cholerae* dataset under the extended Beta-coalescent.

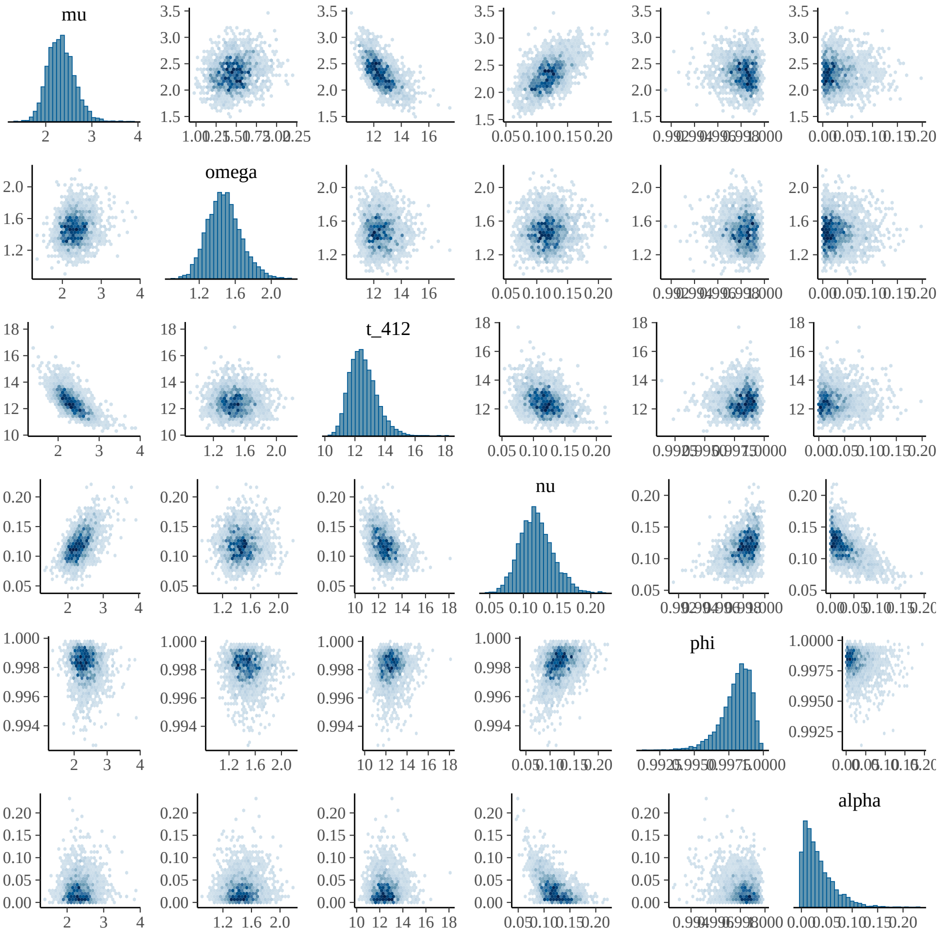


Figure S22: Estimated posterior parameter marginals for the analysis of the *Vibrio cholerae* dataset under extended Beta-coalescent.

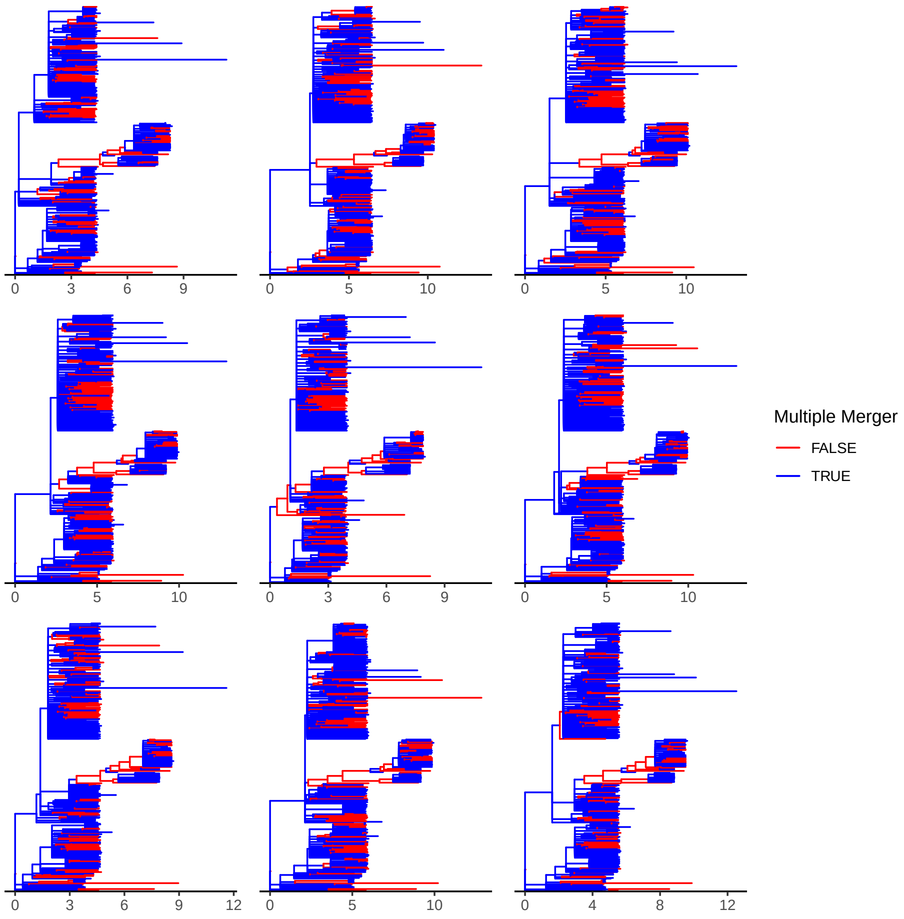


Figure S23: Nine posterior realisations of the underlying genealogy for the analysis of the *Vibrio cholerae* dataset under the extended Beta-coalescent.

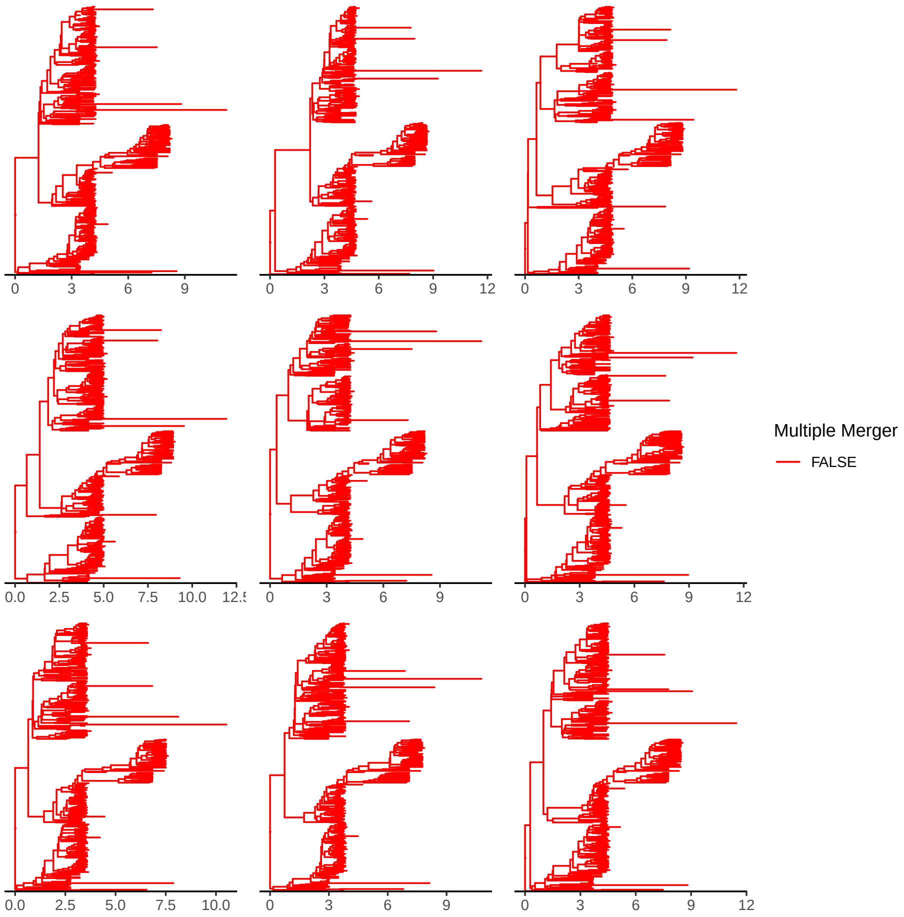


Figure S24: Nine posterior realisations of the underlying genealogy for the analysis of the *Vibrio cholerae* dataset under Kingman's coalescent.

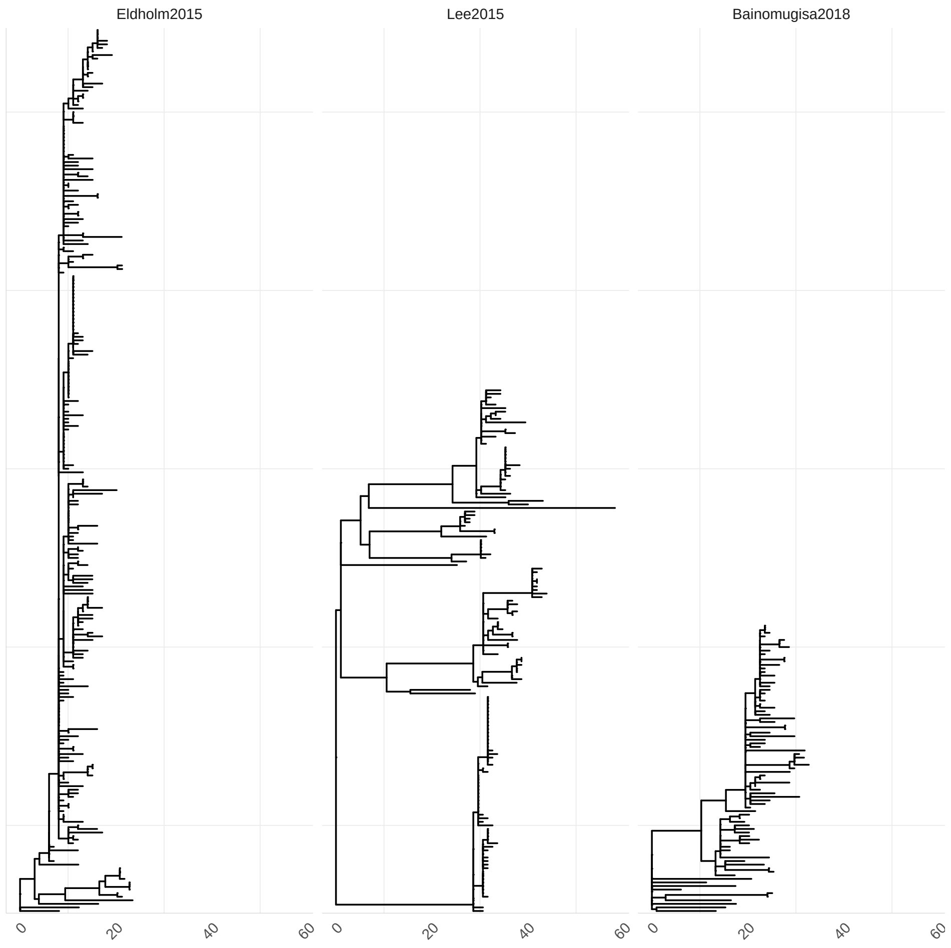


Figure S25: Maximum Likelihood Phylogenies input phylogenies reconstructed using `iqtree -m GTR+G`.

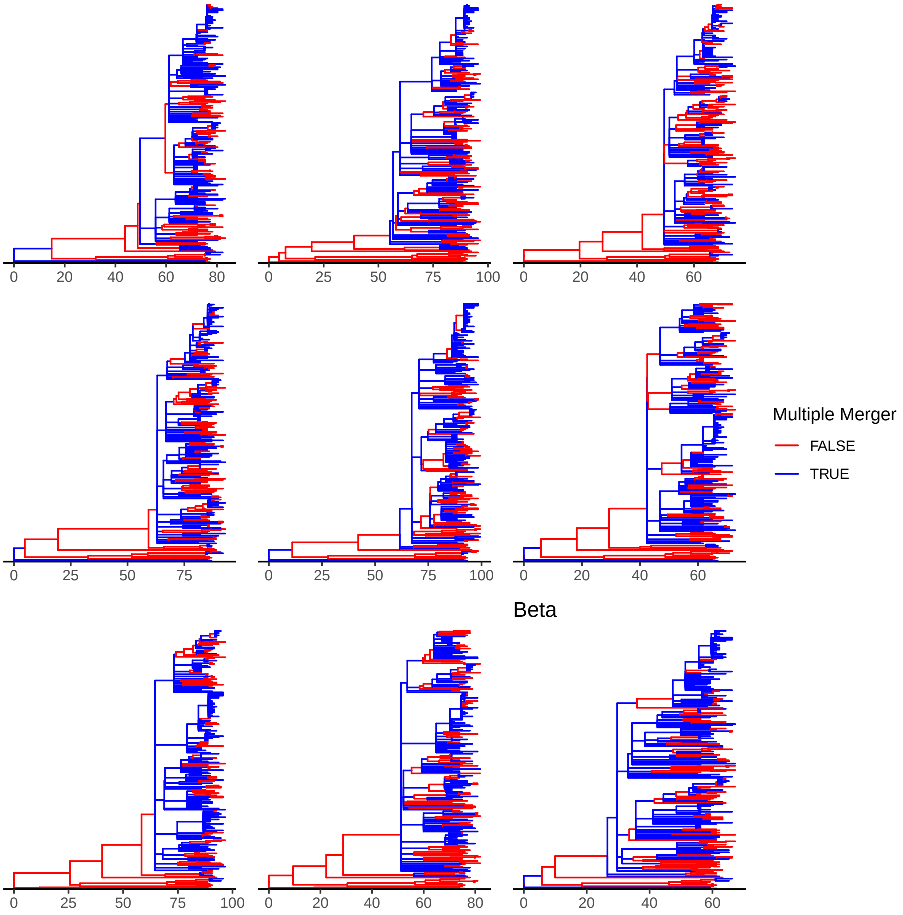


Figure S26: Nine posterior realisations of the underlying genealogy for the analysis of the *Eldholm2015* dataset under the Beta-coalescent.

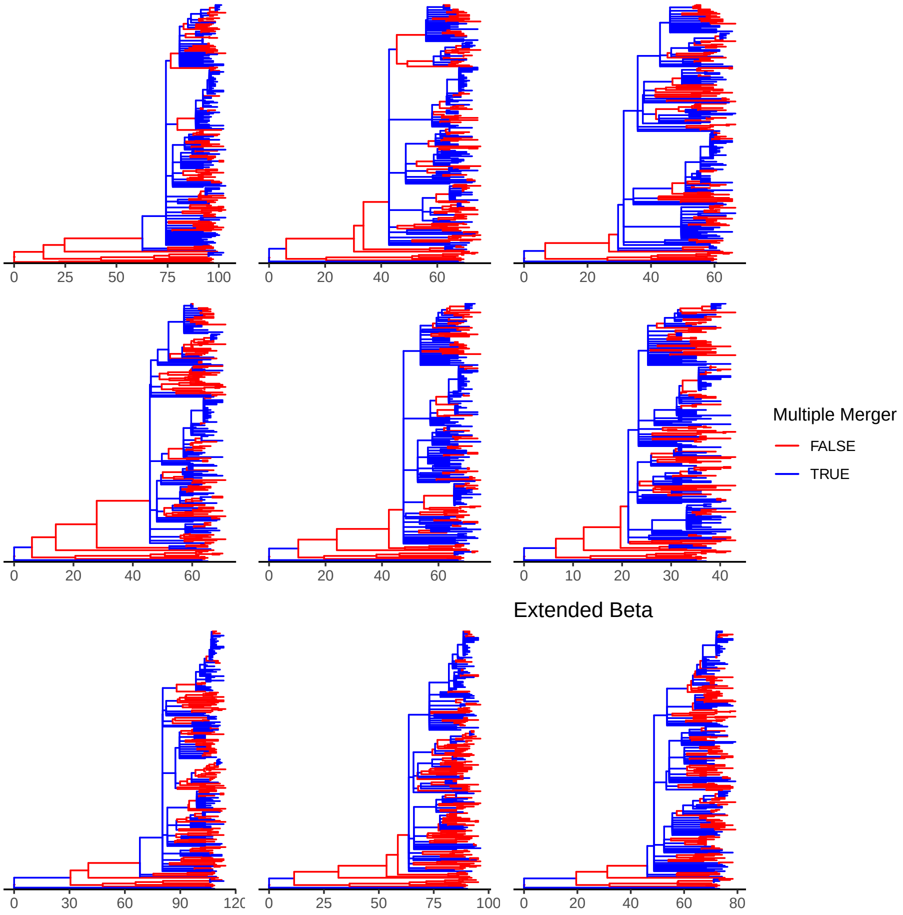


Figure S27: Nine posterior realisations of the underlying genealogy for the analysis of the *Eldholm2015* dataset under the extended Beta-coalescent.

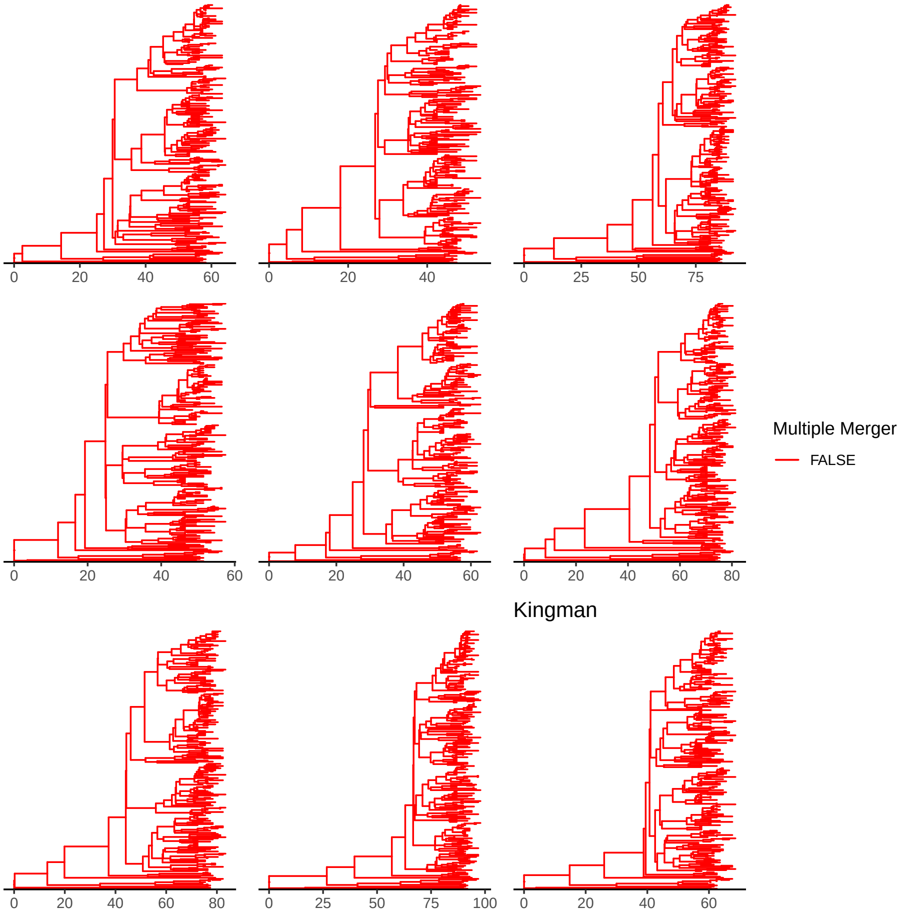


Figure S28: Nine posterior realisations of the underlying genealogy for the analysis of the *Eldholm2015* dataset under Kingman's coalescent.
